# Loss of Kinesin-8 improves the robustness of the acentrosomal spindle

**DOI:** 10.1101/2020.07.29.226506

**Authors:** Alberto Pineda-Santaella, Nazaret Fernández-Castillo, Ángela Sánchez-Gómez, Alfonso Fernández-Álvarez

## Abstract

Chromosome segregation in female meiosis is inherently error-prone, among other reasons, because the acentrosomal spindle assembles and segregates chromosomes without the major microtubule-organizing centres in eukaryotes, the centrosomes, which causes high rate of aneuploidy. The molecular basis underlying formation of acentrosomal spindles is not as well-understood as that of centrosomal spindles and, consequently, strategies to improve spindle robustness are difficult to address. Recently, we noticed during fission yeast meiosis the formation of unexpected microtubules arrays, independent of the spindle pole bodies (yeast centrosome equivalent), with ability to segregate chromosomes. Here, we establish such microtubules formation as bonafide self-assembled spindles that depend on the canonical microtubule crosslinker Ase1/PRC1, share similar structural polarity and harbour the microtubule polymerase Alp14/XMAP215, while being independent of conventional γ-tubulin-mediated nucleation mechanisms. Remarkably, acentrosomal spindle robustness was reinforced by deletion of the Klp6/Kinesin-8, which, consequently, led to a reduced meiotic aneuploidy rate. Our results enlighten the molecular basis of acentrosomal meiosis, a crucial event in understanding gametogenesis.

## Introduction

Meiosis is a specialized type of cell division in which diploid progenitor cells undergo two reductional divisions, named meiosis I (MI) and meiosis II (MII), to produce haploid cells (Griswold and Hunt, 2013). Chromosome segregation during meiotic divisions is mediated by the spindle, that consists of microtubules together with a vast cohort of structural and regulatory proteins. In mitosis and male meiosis, spindle formation is mediated by the centrosomes, the major microtubule-organizing centre in the cell, which localize at the spindle poles (Walczak and Heald, 2008). Centrosomes contain two centrioles, microtubule-derived structures, surrounded by the pericentriolar material, a proteinaceous electron-dense matrix (Bettencourt-Dias and Glover, 2007). In contrast, in female meiosis, centrosomes are purposefully eliminated in oocytes, the gamete progenitor cells, before or during the meiotic divisions (Dumont and Desai, 2012). Generally, oocytes degrade specifically centrioles and maintain the pericentriolar material (Schatten, 1994). Early eukaryotes species of echinoderms (Nakashima and Kato, 2001; Sluder et al., 1993), bivalvia (Longo and Anderson, 1969) and crustaceans (Ruthmann, 1959) enter meiosis with 4 centrioles (2 centrosomes) and gradually expel them out to the degenerating daughter cells in each meiotic division, leaving the mature oocyte with one centriole, which ends up disintegrating. Late eukaryotes are akin to degrade centrosomes in earlier meiotic stages, showing a wide range of differences between species. *Drosophila melanogaster* oocytes initially possess a multicentriolar microtubule-organizing centre but this is fully degraded before the onset of meiotic divisions (Gonzalez et al., 1998; Cooley and Theurkauf, 1994; Colombié et al., 2008; Theurkauf and Hawley, 1992). Similarly, in *Caenorhabditis elegans oocytes* (Kemp et al., 2004; Srayko et al., 2006; Wolff et al., 2016), *Xenopus* extracts (Gard, 1991; Heald et al., 1996; Walczak et al., 1998) and human oocytes (Hertig and Adams, 1967; Sathananthan et al., 2006) centrioles are lost throughout pre-meiotic phases. As a consequence of centriole degradation, the meiotic spindle in these late species must form and segregate chromosomes in the absence of complete, functional centrosomes, being that the reason why this particular type of spindle is known as acentrosomal spindle. Interestingly, mouse oocytes are an exception to this paradigm, because they substitute centrosomes by multiple acentriolar microtubule-organizing centres (Szollosi et al., 1972) that eventually collapse to form the meiotic spindle poles (Kolano et al., 2012; Schuh and Ellenberg, 2007; Clift and Schuh, 2015). The absence of centrosomes in most of female meiosis is one of the most important causes of the high rate of aneuploidy in gametes; in other words, chromosome segregation mediated by acentrosomal spindles is an error-prone process (Holubcova et al., 2015), whose molecular basis is not well-understood due to, among other reasons, the low availability of mammalian oocytes. In this work, we developed a system using the fission yeast *Schizosaccharomyces pombe* to explore the mechanisms behind acentrosomal spindle formation with the aim to improve the robustness of the acentrosomal spindles and, consequently, reduce the impact of aneuploidy during gametogenesis.

Mitotic and meiotic spindles are nucleated in *S*. *pombe* by the spindle pole bodies (SPBs, the centrosome-equivalent in yeast). During interphase, one SPB is sitting on the cytoplasmic side of the nuclear envelope (NE), it duplicates into two SPBs once mitotic and meiotic cell cycle is initiated and then, these get inserted into the partially disassembled portion of NE underneath them (Ding et al., 1997; Bestul et al., 2017) while the rest remains intact (Yoshida and Sazer, 2004); this insertion allows SPBs access the nucleoplasm and start to organize nuclear microtubules into a spindle which eventually segregates chromosomes. Localised NE disassembly and SPB insertion processes are controlled by the interaction of specialised regions of chromosomes, centromeres (in mitosis) and telomeres (in meiosis) and the LINC complex (linker of nucleoskeleton and cytoskeleton) (Tomita and Cooper, 2007; Fennell et al., 2015; Fernández-Álvarez et al., 2016). The LINC complex is composed of the inner nuclear membrane SUN- and outer nuclear membrane KASH-domain proteins, which link centromeres and telomeres with the SPBs (Rothballer et al., 2013). In particular during meiotic prophase, telomeres are gathered beneath the SPB in a chromosomal arrangement called the telomere bouquet (Niwa et al., 2000), being bridged via the telomeric proteins Taz1 and Rap1 and the meiosis-specific Bqt1/Bqt2 dimer to Sad1, the SUN-domain protein in fission yeast (Chikashige et al., 2006). Disruption of telomere-Sad1 interaction, e.g. by deletion of *bqt1*, leads to inhibition of bouquet formation, localised NE disassembly and insertion of SPBs, which remain uninserted distant from the NE; consequently, the absence of the bouquet abolishes SPB-mediated spindle formation (Tomita and Cooper, 2007). Centromeres have the ability to substitute telomeres controlling meiotic SPB insertion into the NE and spindle formation in bouquet-deficient cells due to their residual interaction with Sad1 (Fennell et al., 2015). Interestingly, a combined double point-mutation of *sad1, sad1*.*2* (*sad1*.*T3S*.*S52P*) impairs meiotic prophase centromere-Sad1 interaction, which renders ∼100% of *bqt1*Δ *sad1*.*2* cells unable to insert the SPBs into the NE and form a SPB-mediated spindle (Fernández-Álvarez et al., 2016; Pineda-Santaella and Fernández-Álvarez, 2019). In this scenario, we recently noticed the formation of a nuclear microtubule array structure, with ability to segregate chromosomes (Pineda-Santaella and Fernández-Álvarez, 2019). These findings prompted us to hypothesize that this structure might be a type of self-assembled spindle similar to that of mammalian acentrosomal meiosis. In this study, we establish that the microtubule organization observed in the absence of SPB insertion is a proper acentrosomal spindle and we convey a further molecular characterization of this type of self-assembled spindle in fission yeast with the aim to shed light on the molecular basis of acentrosomal spindle structure and function. Finally, we design a strategy to increase the robustness of the self-assembled spindles, which allowed to improve the efficiency of chromosome segregation.

## Results

### Self-assembled spindles formation and polarization is independent of LINC-SPB complex

A key question in our recent observations of the formation of an array of nuclear microtubules in the absence of the SPB insertion is whether these are a type of self-assembled spindle. Although we observed that the formation of the array is independent of the SPB (Pineda-Santaella and Fernández-Álvarez, 2019), other SPB-associated elements placed in the NE might be mediating its formation and/or polarization. With this possibility in mind, we decided to follow the behaviour of the LINC complex that, in fission yeast mitosis and meiosis, is permanently associated to the inner part of the SPB (Hagan and Yanagida, 1995; Rothballer et al., 2013). In particular, we observed SUN-domain protein Sad1, the most internal component of the LINC complex, facing towards the nucleoplasmic environment. To do this, we endogenously tagged Sad1 to visualise by live fluorescence microscopy its location under conditions of absence of SPB insertion in meiosis, which requires the double mutation *bqt1Δ sad1*.*2* (Fernández-Álvarez et al., 2016). We also compared the behaviour of Sad1-GFP and Sad1.2-GFP throughout meiosis in *bqt1*^*+*^ cells (showing SPB-mediated spindles) as a control to verify that *sad1*.*2* mutations do not affect Sad1 protein location (Fernández-Álvarez et al., 2016) (Fig. 1A 25’ & 1B 30’, yellow arrowheads). Analysis of *bqt1Δ sad1*.*2-GFP* meiosis showed the formation of an array of polarized microtubules organized around chromosomes (self-assembled spindles) with capability to segregate them (Fig. 1C, 20’ to 40’). Remarkably, Sad1.2 location at the tips of the microtubules array is excluded in contrast to the SPB-mediated spindles (compare 1A or 1B *vs* 1C 15’, yellow arrowhead), indicating that polarization of the microtubule array is independent of the SUN-domain protein location. These observations together with our previous data showing a similar behaviour of the integral SPB components Sid4, Cut12 and Pcp1 to those observed in *bqt1Δ sad1*.*2-GFP* meiosis (Pineda-Santaella and Fernández-Álvarez, 2019) indicate that the array of microtubules is a type of self-assembled spindle whose formation and polarization are not mediated by the localization of the LINC complex nor the SPB (Fig. 1D).

**FIGURE 1.**
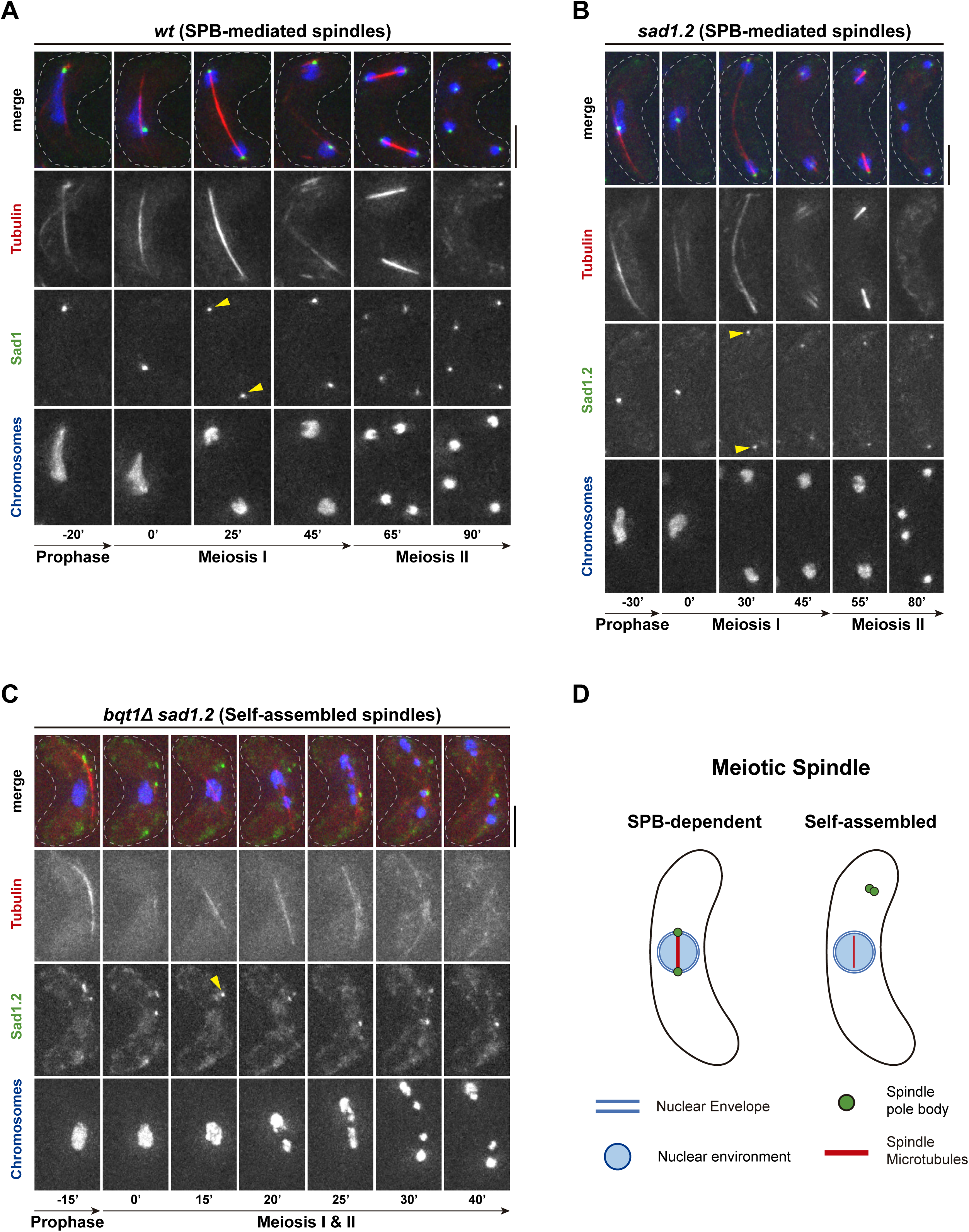
Spindle self-assembly occurs independently of the LINC complex. **(A)** In a wild-type meiosis, Sad1, most inner component of the LINC complex in direct contact with the spindle pole body, localizes at the poles of the SPB-mediated meiotic spindle in MI (25’) and MII (65’). **(B)** The double point mutation T3S, S52P of the mutant allele *sad1*.*2* of *sad1* do not alter the normal localization of Sad1 protein in meiosis (compare A 25’ and B 30’, yellow arrowheads). **(C)** Sad1.2 does not localize at the poles of self-assembled spindles (yellow arrowhead), which means that spindle self-assembly is not mediated by the LINC complex, as it happens for SPB-mediated meiotic spindles. **(D)** Schematic comparison between meiotic SPB-mediated and self-assembled spindles. Sad1/Sad1.2: Sad1/Sad1.2-GFP; For this and the rest of the figures, microtubules are visualized via *mCherry-atb2* (α2 tubulin subunit) and chromosomes via *hht1-CFP* (histone H3) as well as scale bars correspond to 5 µm and timestamps are referred to the onset of spindle formation.

### Ase1/PRC1 is an essential structural component of meiotic self-assembled spindles

To further substantiate the idea that self-assembled spindles in *bqt1Δ sad1*.*2* meiosis share the canonical features of a proper spindle, we carried out a characterization of the elements controlling its dynamics. We previously observed that the microtubule crosslinker protein Ase1/PRC1 localizes to the body of self-assembled spindles in very similar patterns to those seen for SPB-mediated spindles, including at the spindle midzone (Pineda-Santaella and Fernández-Álvarez, 2019). Ase1 location at the midzone in self-assembled spindles suggests that this protein might be part of the structure and be involved in its formation and extension. In order to unclose the functional relevance of Ase1 for self-assembled spindles behaviour, the effect of *ase1* deletion was analysed in meiosis. Stages of *ase1*^*+*^ SPB-mediated spindles comprises nucleation (Fig. 2 A, 0’), assembly (Fig. 2 A, 0’ to 25’), elongation (Fig. 2 A, 25’ to 30’) and disassembly, which consists of a dismantling of the whole spindle body (Fig. 2 A, 30’ to 40’). Consistently, we observed similar stages for *ase1*^*+*^ self-assembled spindles comprising formation (Fig. 2 B, 0’ to 15’), elongation (Fig. 2 B, 15’ to 35’), and disassembly (Fig. 2 B, 35’ to 35’ to 45’). Interestingly, *ase1Δ* MI SPB-mediated spindles exhibit normal nucleation and assembly (Fig. 2 C, 0’ to 50’), but, in contrast to a *ase1*^*+*^ disassembly, *ase1Δ* cells show a discrete breakage at the spindle midzone during elongation from which spindle microtubules shrink, leading to spindle dissolution (Fig. 2 C 55’ to 70’, yellow arrowheads). This phenotype can be also observed during MII (Fig. 2 C, 105’, yellow arrowhead). Strikingly, *ase1Δ* self-assembled spindles displayed similar behaviour compared to that of *ase1Δ* SPB-mediated spindles (Fig. 2 D 50’ to 60’, arrowheads). Congruently, for both SPB-mediated and self-assembled spindles, the maximum spindle length is significantly shorter upon *ase1* deletion (Fig. 2 E), indicating that spindle breakage takes place in a premature manner during elongation, whereas in *ase1*^*+*^ settings the spindle keeps elongating until normal disassembly occurs with a longer length. These results establish that Ase1/PRC1 is essential for the maintenance of the structural integrity of self-assembled spindles in meiosis.

**FIGURE 2.**
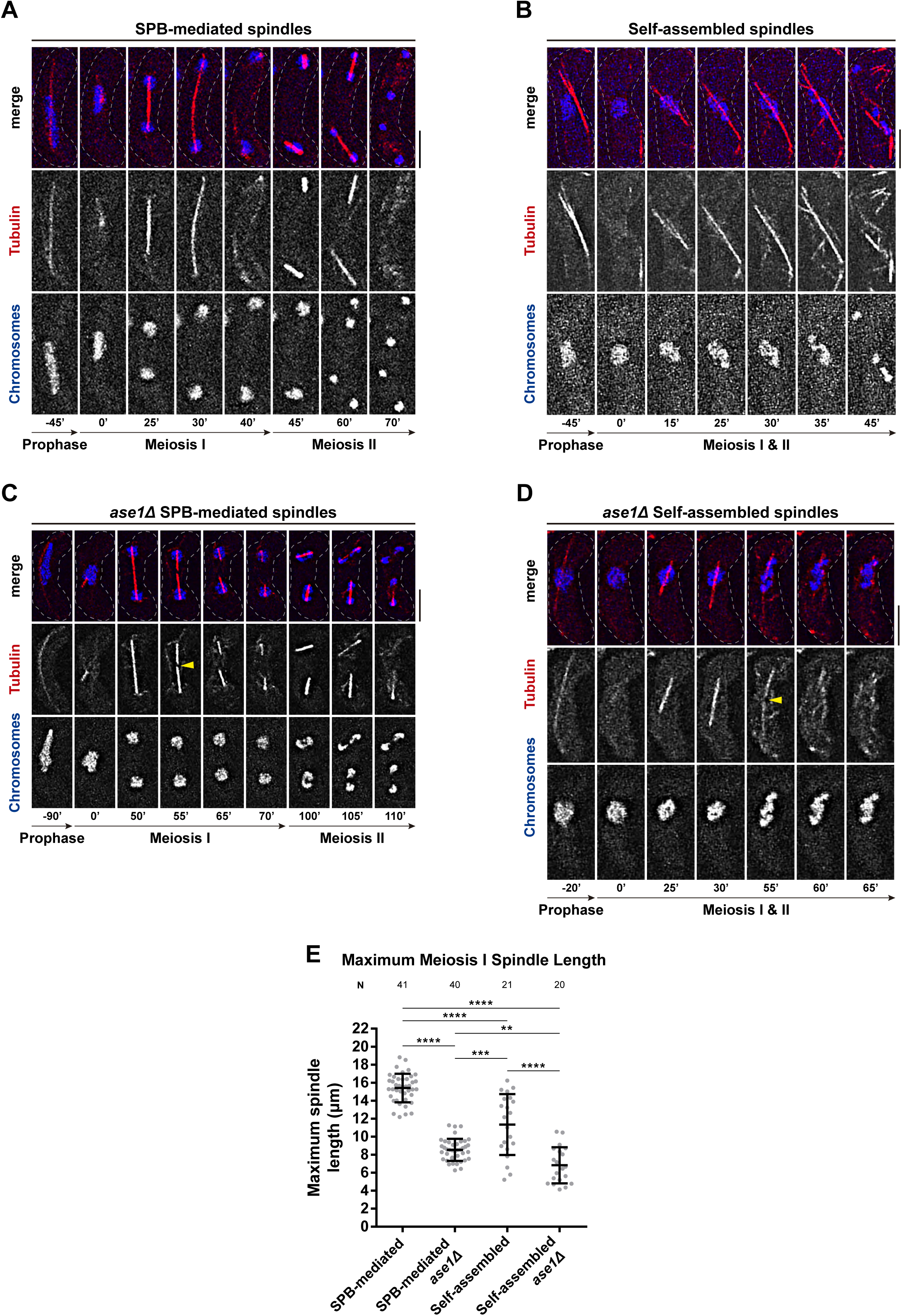
Microtubule crosslinker Ase1/PRC1 is an essential component of self-assembled spindles. **(A)** In a wild-type setting, SPB-mediated spindles elongate up to a maximum length and then the whole spindle body is disassembled simultaneously in MI (30’ to 40’) as well as MII (60’ to 70’). **(B)** Similar to SPB-mediated spindles, self-assembled spindles elongate up to a maximum length at which they disassemble as a whole. **(C)** Upon *ase1* deletion, the wildtype MI spindle suffers a discrete breakage at the midzone during elongation (55’, yellow arrowhead) and spindle microtubules shrink from that breakage point (65’ to 70’), similarly happening for MII spindles (105’, top spindle). **(D)** Upon *ase1* deletion, self-assembled spindles suffer a discrete breakage at the midzone (55’, yellow arrowhead). **(E)** Quantification of maximum spindle length shows that *ase1* deletion shortens the maximum spindle length in SPB-mediated and self-assembled backgrounds. t-test with Welch’s correction: ****: p < 0.0001, ***: p = 0.0013, **: p = 0.0017.

### Microtubules arrangement in self-assembled spindles shows minus ends at the poles

We have shown that self-assembled spindles are organised around chromosomes and grow with a polarity that is not determined by the SPB-LINC complex; for this reason, we wanted to establish if these spindles are characterised by a normal or inverted polarity. The fact that Ase1 is required for the dynamics of self-assembled spindles (Figure 2) together with its location at spindle midzone (Pineda-Santaella and Fernández-Álvarez, 2019) strongly suggest the existence of a central zone composed of overlapping antiparallel microtubules in self-assembled spindles. Ase1 location at the midzone can be explained by two different plausible spindle microtubules configurations: i) *plus-end* microtubules at the midzone and *minusends* at the tips (SPB-mediated spindle polarity) or, the opposite, ii) *minus-end* microtubules at the midzone and *plus-ends* at the tips (similar to that of interphase microtubules arrays). To establish the polarity of self-assembled spindles, we analysed the localization of a GFP-tagged version of Pkl1/Kinesin-14 (Pkl1-GFP), a kinesin that specifically tracks spindle microtubules *minus-ends* (Pidoux et al., 1996; Yukawa et al., 2015). As reported for SPB-mediated mitotic spindles, Pkl1-GFP localizes to the spindle poles in MI (Fig. 3 A, 20’ to 35’) and MII (Fig. 3 A, 80’ to 90’). Remarkably, Pkl1-GFP also localizes to the poles of self-assembled spindles (Fig. 3 B, 20’ to 30’), indicating that the tips of these spindles are composed of microtubules *minus-ends*. Hence, structural polarity of self-assembled spindles obeys the microtubules arrangement with *minus* ends at the poles and *plus* ends at the midzone, resembling that of SPB-mediated mitotic and meiotic spindles.

**FIGURE 3.**
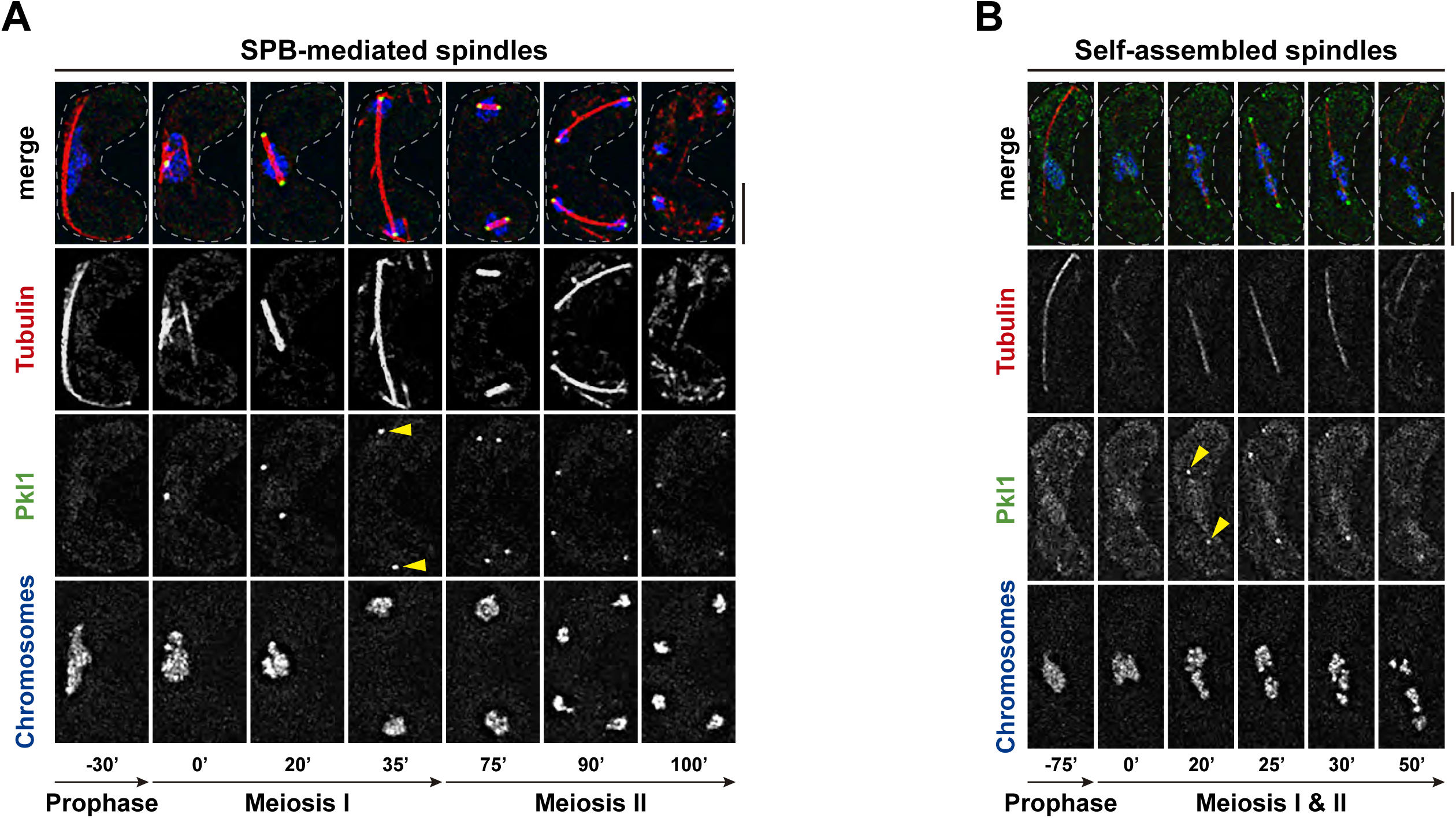
Poles of self-assembled spindles correspond to microtubules minus-ends. **(A)** In a wildtype meiosis, Pkl1, a kinesin motor directed to microtubule minus-ends, localizes to the poles of MI (20’ to 35’) and MII (80’ to 90’) SPB-mediated spindles. **(B)** Similarly, Pkl1 also localizes to the poles of self-assembled spindles (20’ to 30’, yellow arrowheads), showing that these poles correspond to the minus-ends of microtubules that build up the spindle. Pkl1: Pkl1-GFP.

### F-actin network is dispensable for self-assembly spindle formation

To further characterise the formation and dynamics of self-assembled spindles, we wanted to explore the possible role of other cytoskeleton components. In this context, F-actin plays a role in chromosome segregation in oocytes (Mogessie and Schuh, 2017). In fission yeast, F-actin network is important for proper spindle orientation in mitosis (Gachet et al., 2001, Gachet2006), but its role for meiotic spindle formation and behaviour has not been yet disclosed. Given this uncertainty, we decided to explore whether F-actin network is important for formation and dynamics of SPB-dependent and self-assembled spindles during fission yeast meiosis. For this purpose, F-actin network was disrupted using the actin-depolymerizing drug Latrunculin A (LatA), specifically during meiosis (see Methods) and then spindle behaviour was analysed. In order to follow F-actin network, we used the fluorescent label Life-Act (Riedl et al., 2008) and monitored it together with chromosomes and microtubules. In the absence of LatA, F-actin can be observed as cables (Fig. S1 A, red arrowheads) and numerous patches which fluctuate throughout the cell body during prophase and MI (Fig. S1 A, −100’ to 35’), then it collapses and assembles into four meiotic actin rings surrounding post-MI nuclei during MII (Fig. S1 A, 85’ to 90’) and these eventually contract (Fig. S1 A, 95’) and disassemble, congruent with previous description in meiosis (Yan and Balasubramanian, 2012). In this setting, SPB-dependent spindles display normal assembly, elongation and disassembly dynamics along with symmetrical segregation of chromosomes into four equal-size masses (Fig. S1 A, 0’ to 95’). Under addition of 4 µM LatA, F-actin patches signal is significantly reduced (Fig. S1 B and C, yellow rectangles, ‘actin fading’ in down panels) (Fig. S1 C), although it is recovered later on as meiosis progresses (Fig. S1 B and C, yellow rectangles, ‘actin recovery’ in bottom panels). Additionally, actin cables are almost completely absent after the treatment (compare Fig. S1 A, −100’ and −75’ vs Fig. S1 B). Hence, we confirm that LatA is bioactive in our experimental conditions, achieving a partial F-actin depolymerisation upon drug addition in meiosis. Noteworthy, the effect of the depolymerizing activity of the drug extends throughout the whole meiosis as shown by the fact that actin rings, taking place after actin patches recovery, are also affected (Fig. S1 B 145’ and C 180’).

Next, we analysed the potential role of F-actin in self-assembled spindle formation and dynamics. In the absence of LatA, the behaviour of F-actin patches (Fig. 4 A, −95’ to 0’) and cables (Fig. 4 A, −95’ to −50’, red arrowheads) is comparable to that seen for SPB-mediated spindles, although no normal actin rings are observed: around 9% of cells show defective actin rings while 91% show no actin rings at all, revealing that actin rings defects is a consequence of losing the SPBs insertion. Upon 4 µM LatA treatment, F-actin is depolymerised (Fig. 4 B, yellow rectangles, “F-actin fading” in down panels and 4 C; Fig. 4D) as we expected. Noteworthy, formation and dynamics of self-assembled spindles were similar to those observed in the absence of LatA (Fig. 4 B, 0’ to 35’), indicating that F-actin disruption does not have an impact on self-assembled spindle formation (Fig. 4 F). Altogether, these findings illustrate that meiotic F-actin network does not play a key role in neither the formation nor the dynamics of self-assembled spindles.

**FIGURE 4.**
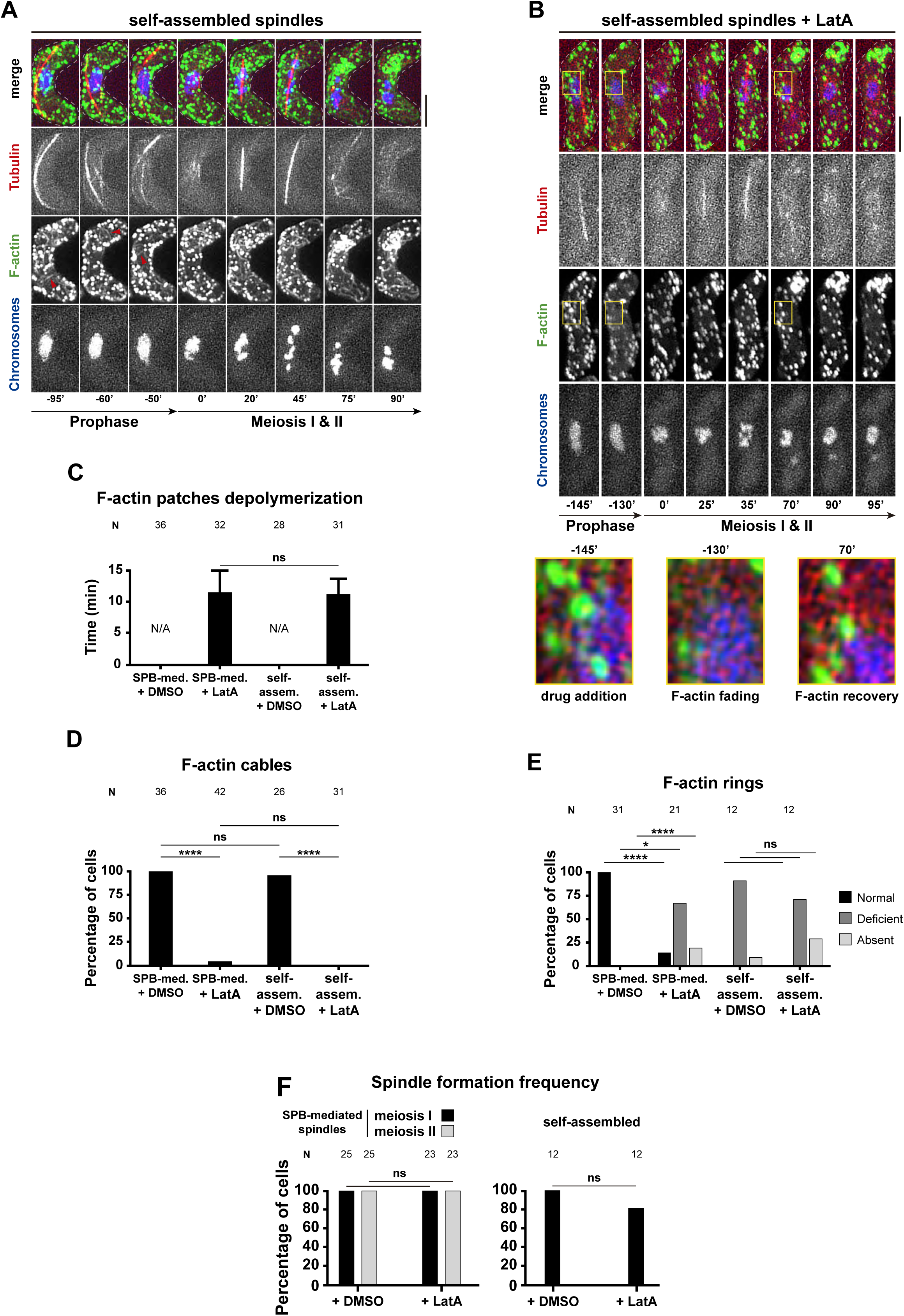
Self-assembled spindles formation and dynamics are independent of the F-actin network in meiosis. **(A)** In a self-assembled spindle background, in meiotic prophase, microtubules oscillate disconnected from the nucleus, which stays immobile at the cell centre (−95’ to −50’). After prophase, self-assembled spindles form (0’ to 20’), segregate chromosomes (20’ to 45’) and disassemble (75’ to 90’). Unaltered, F-actin network display disperse patches and thin actin cables during prophase (−95’ to −50’) but in MII shows aberrant aggregates. **(B)** Upon addition of actin-depolymerizing drug Latrunculin A (LatA), actin cables are no longer observed, F-actin patches signal is reduced (zoomed areas indicated with rectangles) and actin rings are not observed. In spite of F-actin disruption, self-assembled spindles still form with similar morphology and dynamics (25’ to 35’) and are able to segregate chromosomes (35’ to 70’), as in the absence of drug. **(C)** Quantification of time needed to start seeing F-actin depolymerisation from the moment of drug addition. F-actin depolymerisation is not detected in controls with only the solvent used to dissolve the drug (DMSO). No significant difference is observed in depolymerisation times between SPB-mediated and self-assembled spindles backgrounds (“ns”, t-test: p > 0.05). **(D)** Upon LatA treatment, percentage of cells displaying actin cables significantly decreases in both SPB-dependent and self-assembled spindles backgrounds (Fisher exact test, ****: p < 0.0001 for both cases), observing no significant difference between LatA treated cases (Fisher exact test, “ns”: p > 0.05). **(E)** Quantification of actin rings behaviour shows that LatA addition leads to severe alteration of actin rings morphology and formation in the SPB-mediated spindles background (Fisher exact test, ****: p < 0.0001, *: p = 0.0221), while no significant effect is observed in the self-assembled spindle background (Fisher exact test, “ns”: p > 0.05). **(F)** Spindle formation frequency shows no significant difference between absence and presence of LatA for SPB-dependent and self-assembled spindles (Fisher exact test, “ns”: p > 0.05, in both cases). F-actin: Lifeact-GFP.

### SPB-associated γ-tubulin complex is not involved in self-assembled spindle nucleation

After completion of meiotic prophase, cytoplasmic astral oscillating microtubules dissolve just before the formation of SPB-mediated MI and MII spindles. Nucleation of microtubules is carried out by a macromolecular protein complex called the γ-tubulin complex, which serves as a structural template for priming the *de novo* synthesis of microtubule filaments (Braunfeld et al., 2002). At the time of spindle nucleation, this complex actively targets to the nuclear side of the already inserted SPBs, nucleating microtubules that are then elongated via polymerization, projecting from the SPBs towards the nucleoplasm (Vardy, 2000; Bestul et al., 2017). In the case of self-assembled spindles, the SPBs are not inserted into the NE, thus, the possible role of γ-tubulin complex is unclear. To investigate whether self-assembled spindles nucleation depends on this mechanism, a GFP-tagged version of Alp4 (Alp4-GFP), an essential component of the γ-tubulin complex (Vardy, 2000), was used as a proxy to monitor the localization of the complex relative to self-assembled spindles. In a control scenario of SPB-mediated spindles, one Alp4-GFP dot is observed during meiotic prophase to localize at the leading edge of the astral microtubules structure, following its oscillating movement (Fig. 5 A, −25’). After prophase, one dot of Alp4-GFP co-localizes with the microtubules focus from which the spindle emerges at the time of its formation (Fig. 5 A, 0’), consistent with the idea of γ-tubulin complex recruitment to the SPBs (Horio et al., 1991; Masuda et al., 2013). Later, concomitantly with MI spindle assembly and elongation, the original Alp4-GFP dot splits into two separate dots which perfectly co-localize with the spindle poles (Fig. 5 A, 0’ to 35’). After MI spindle disassembly, the same behaviour is observed for MII SPB-mediated spindles (Fig. 5 A, 45’ to 100’). Eventually, after MII spindles disassembly, each of the four resulting Alp4-GFP dots remain localizing to each of the four resulting chromosomes masses (Fig. 5, A 115’ to 120’). For self-assembled spindles, Alp4-GFP equally follows the leading edge of the oscillating astral microtubule structure, with the difference that the nucleus does not follow the oscillations, unlike the SPB-mediated spindle background (Fig. 5 B, −85’ to −80’). Strikingly, after prophase, while the self-assembled spindle forms within the chromosomal environment, the Alp4-GFP dot stays dislodged, far from the self-assembled spindle and chromosomes (Fig. 5 B, 0’ to 45’, yellow arrowheads). Furthermore, in a subset of cells, the original Alp4-GFP dot is observed to split up to four dots (Fig. 5 B, 75’, yellow asterisk) together with the appearance of microtubule filaments in their vicinity (Fig. 5 B, 45’ to 75’, orange asterisks), pointing that these dots correspond to γ-tubulin complex associated to the uninserted SPBs. Thus, this dissociation between the self-assembled spindles and the Alp4 dots suggests that the nucleation of self-assembled spindles is independent of the conventional nucleation mechanism driven by the SPB-associated γ-tubulin complex, contrary to the case of SPB-mediated spindles.

**FIGURE 5.**
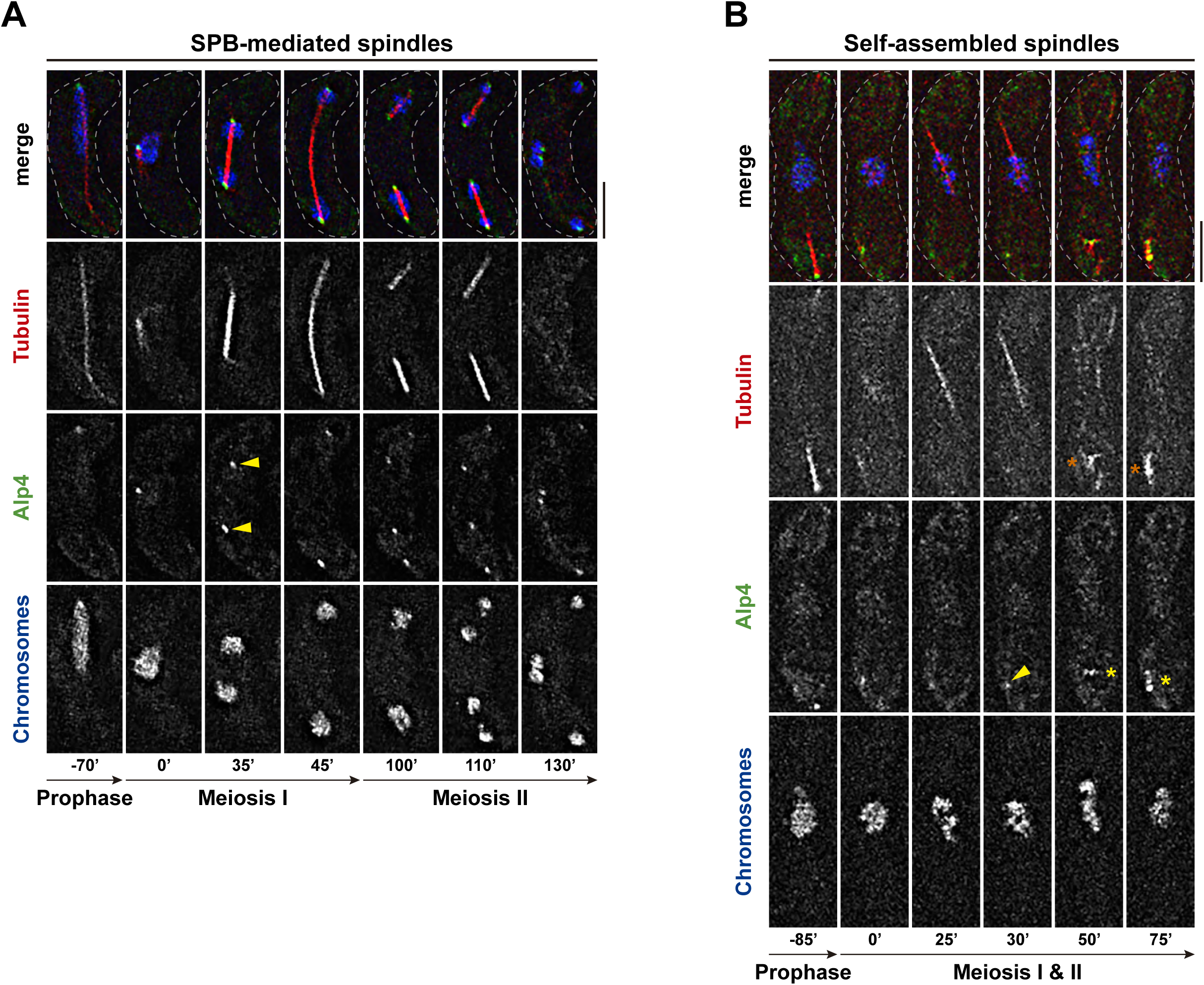
Self-assembled spindles formation is independent of SPB-associated γ-tubulin complex. **(A)** In the SPB-mediated spindles background, Alp4 localizes as one dot at the forefront of the oscillating microtubules in the meiotic prophase (−25’) and later localizes as one dot at each pole of MI (35’ to 45’) and MII (100’ to 115’) spindles. **(B)** In the self-assembled spindle background, the behaviour of Alp4 in prophase is similar, but later, it stays as a dot far away from the self-assembled spindle while it forms (0’ to 45’, yellow arrowheads). After spindle disassembly, microtubules start to form around this Alp4 dot (45’ to 75’, orange asterisks), which appear to split into several dots (75’, yellow asterisk). Alp4: Alp4-GFP.

### Microtubule polymerase Alp14/XMAP215 accumulates around self-assembled spindles

Our previous results suggest that the γ-tubulin complex is not implicated in self-assembled spindle formation via the conventional nucleation mechanism. For this reason, we decided to investigate the possible role of other factors involved in microtubule nucleation and polymerization in controlling the formation of self-assembled spindles. In fission yeast, the microtubule polymerase XMAP215 family is represented by the well-characterised Alp14 protein (Al-Bassam et al., 2012). Alp14 participates in the nucleation of interphase microtubule arrays (Flor-Parra et al., 2018) as well as the formation of the mitotic spindle through its microtubule polymerization activity, contributing to SPBs separation and correct bipolar spindle assembly (Yukawa et al., 2017). However, knowledge on its role during meiosis has not been well-established yet. With the aim to explore the behaviour of Alp14 during fission yeast meiosis and, in particular, in a self-assembled spindle scenario, a GFP-tagged version of Alp14 (Alp14-GFP) was monitored throughout meiosis. For SPB-mediated spindles, during prophase, Alp14-GFP localizes to oscillating astral microtubules accumulating at the forefront of this structure (Fig. 6 A −45’, B −25’, yellow arrowheads). At the onset of SPB-mediated spindle formation, Alp14-GFP colocalizes with the tubulin focus from which the spindle emerges (Fig. 6 A, 0’), suggesting a role in spindle microtubule nucleation. Once the spindle forms, Alp14 shows several localization patterns: i) first, as dots along the body of early short MI spindles (Fig. 6 A, 15’), as reported for the mitotic spindle (Garcia, 2001; Al-Bassam et al., 2012); ii) second, to spindle poles (yellow arrowheads) and midzone (asterisk) of late elongated MI spindles (Fig. 6 A, 30’) and iii) third, to the whole body of MII spindles in a continuous manner (Fig. 6 A, 85’). Noteworthy, Alp14-GFP also localizes to elongated self-assembled spindles, concentrating at spindle poles (Fig. 6 B, 15’ to 35’, yellow arrowheads), similar to the behaviour observed in SPB-mediated spindles. However, due to the more dispersed signal of Alp14-GFP around self-assembled spindles, it is difficult to discern its precise localization at the onset of spindle formation (compare Fig. 6 A 0’ vs B 0’) as well as at the midzone of late spindles (compare Fig. 6 A 15’ vs B 15’). Altogether, these findings suggest that Alp14 is involved in the formation and dynamics of self-assembled spindles in a similar manner than in the SPB-mediated spindles.

**FIGURE 6.**
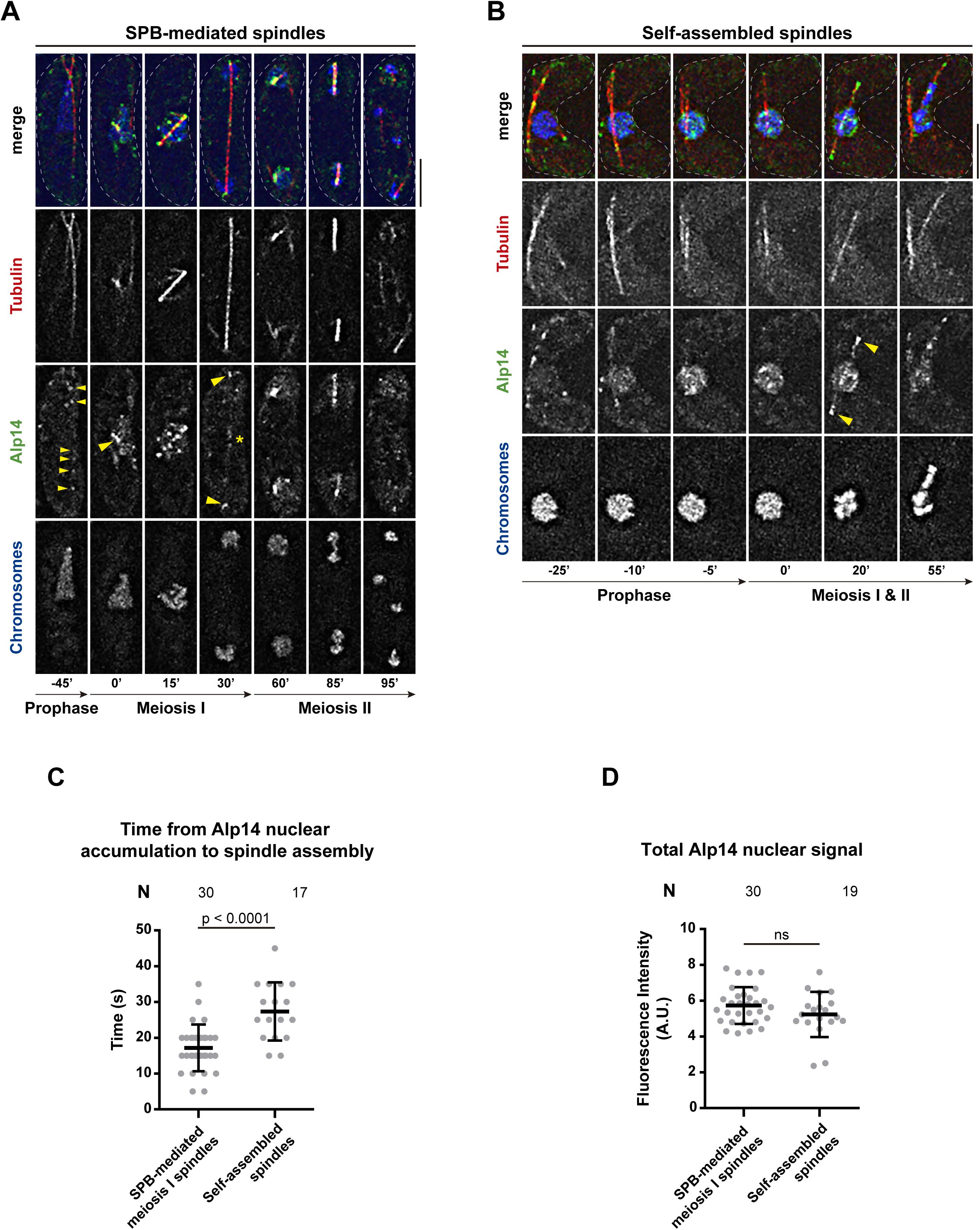
Alp14/XMAP215 microtubule polymerase localizes to self-assembled spindles. **(A)** For SPB-mediated spindles, Alp14 is imported into the nucleus prior to spindle formation (−5’ to 0’), accumulates at the body of MI and II short spindles (15’ and 85’, respectively) and localizes to the midzone (yellow asterisk) and poles (yellow arrowheads) of MI long spindles (30’). **(B)** In a self-assembled spindle background, Alp14 is also imported into the nucleus before spindle formation (−10’ to 0’) and clearly localizes to self-assembled spindles extremes (15’ to 35’, yellow arrowheads), although localization at spindle formation onset and spindle midzone cannot be clearly appreciated due to disperse nucleoplasmic signal of the protein. **(C)** Quantification of the time between Alp14 nuclear accumulation and onset of spindle formation shows a significantly longer time for self-assembled spindle background (t-test, p < 0.0001). Bars: mean and SD. **(D)** Quantification of maximum total nuclear fluorescence signal of Alp14 shows no significant difference between SPB-mediated and self-assembled spindles backgrounds (t-test, “ns”: p > 0.05). Bars: mean and SD. Alp14: Alp14-GFP.

To further characterize the nuclear signal of Alp14-GFP during the self-assembled spindle formation, we analysed Alp14-GFP nucleoplasmic fluorescence intensity dynamics throughout meiosis. We observed that Alp14-GFP accumulates in the nucleus at the end of prophase, noticed by an increasing signal co-localizing with chromosomes, prior to spindle formation (Fig. 6 A -5’, B -10’). Interestingly, the time interval between the initiation of Alp14-GFP nuclear accumulation and spindle formation onset is significantly longer for self-assembled spindles than for SPB-mediated spindles (Fig. 6 C). To check whether this difference was consequence of abnormal Alp14 dynamics, we estimated the amount of nuclear Alp14 in this time window by measuring Alp14-GFP total intensity within the chromosomal environment (see Methods). This quantification showed no significant difference between the maximum levels of nuclear Alp14-GFP of both settings (Fig. 6 D), suggesting that pre-meiotic Alp14 nuclear accumulation remains normal, in favour of the alternative that there is a delay in self-assembled spindle formation, consistent with previously observed defective spindle biophysics (Pineda-Santaella and Fernández-Álvarez, 2019).

### The robustness of self-assembled spindles can be improved by the loss of Kinesin-8

An important characteristic of self-assembled spindles is their low robustness and the consequent high rate of chromosome segregation defects. Specifically, the weakness of the self-assembled spindle structure is more notorious during the second meiotic divisions, with the majority of meiocytes finishing meiosis without forming MII spindles. These defects make the study of the molecular mechanism behind SPB-independent spindle formation difficult, especially in meiosis II. Therefore, we tried to improve the robustness of self-assembled spindles to better get insight into the molecular basis of acentrosomal meiosis. For this purpose, we designed two different strategies: i) the increment of Cls1/CLASP-1 protein levels and, ii) the loss of the Klp6/Kinesin-8. The role of Cls1 is to bundle the overlapping microtubules, regulating the dynamics of interphase microtubule arrays and stabilizing mitotic spindle microtubules (Grallert et al., 2006; Bratman and Chang, 2007; Ebina et al., 2019). Given its role with spindle microtubules, we over-produced Cls1 expecting to increase microtubule stabilization and/or enhance incorporation of more microtubule fibres into the spindle body. On the other hand, we carried out the deletion of Klp6/Kinesin-8. Klp5 and Klp6, the representatives of Kinesin-8 family in fission yeast, are motor proteins which form a heterodimer Klp5/6 involved in regulating microtubule dynamics in interphase and mitosis through diverse activities (Unsworth et al., 2008; Gergely et al., 2016; West and McIntosh, 2008), such as microtubule destabilization. Indeed, deletion of either one leads to depolymerisation-resistant interphase microtubules (Garcia et al., 2002) and promotes spindle microtubules polymerization (Pinder et al., 2019). To assess the possible improvement of the spindle, we quantified the effect of each strategy according to three parameters: i) spindle formation frequency, ii) maximum spindle length as readout of structural strength and iii) chromosome segregation efficiency. Of these two alternatives, we continued to study the deletion of Klp6 because such strategy is technically easier controllable than Cls1 over-expression.

The absence of Klp6 does not produce major defects in MI and MII SPB-mediated spindle formation but *klp6Δ* cells display chromosome congression defects both during mitosis and meiosis, with chromosome masses remaining asymmetrically positioned along the spindle axis before segregation (Fig. 7 B, 20’ arrowhead, 70’ top arrowhead) as well as lagging chromosomes (Fig. 7 B, 70’ bottom arrowhead) (Pinder et al., 2019; Syrovatkina et al., 2013). Remarkably, in the case of self-assembled spindles, deletion of Klp6 rendered MI and MII self-assembled spindles thicker and brighter respect to *klp6*^*+*^ (compare Fig. 7 C 30’ vs D 20’), suggesting an increase of the robustness of the spindle. To better substantiate this observation, we quantified spindle formation frequency, showing that deletion of *klp6* increases the percentage of cells harbouring MII self-assembled spindles from 42% to 67% (respect to the total of cells performing MI, n=55 and n=61, respectively), thus promoting spindle formation and progression to MII. Conversely, there is no significant difference between *klp6*^*+*^ and *klp6Δ* MII SPB-dependent spindles, meaning that Klp6 absence does not significantly affect SPB-dependent meiosis progression (Fig. 7 E). Moreover, we studied the effect on spindle structure by measuring the maximum length of MI spindles. Interestingly, the length of self-assembled spindles (∼8 µm) enlarges upon *klp6* deletion up to normal levels of *klp6*^*+*^ (∼13 µm) and *klp6Δ* SPB-dependent spindles (∼14 µm) (Fig. 7 F). This indicates that deletion of *klp6* is able to strengthen self-assembled spindle structure.

**FIGURE 7.**
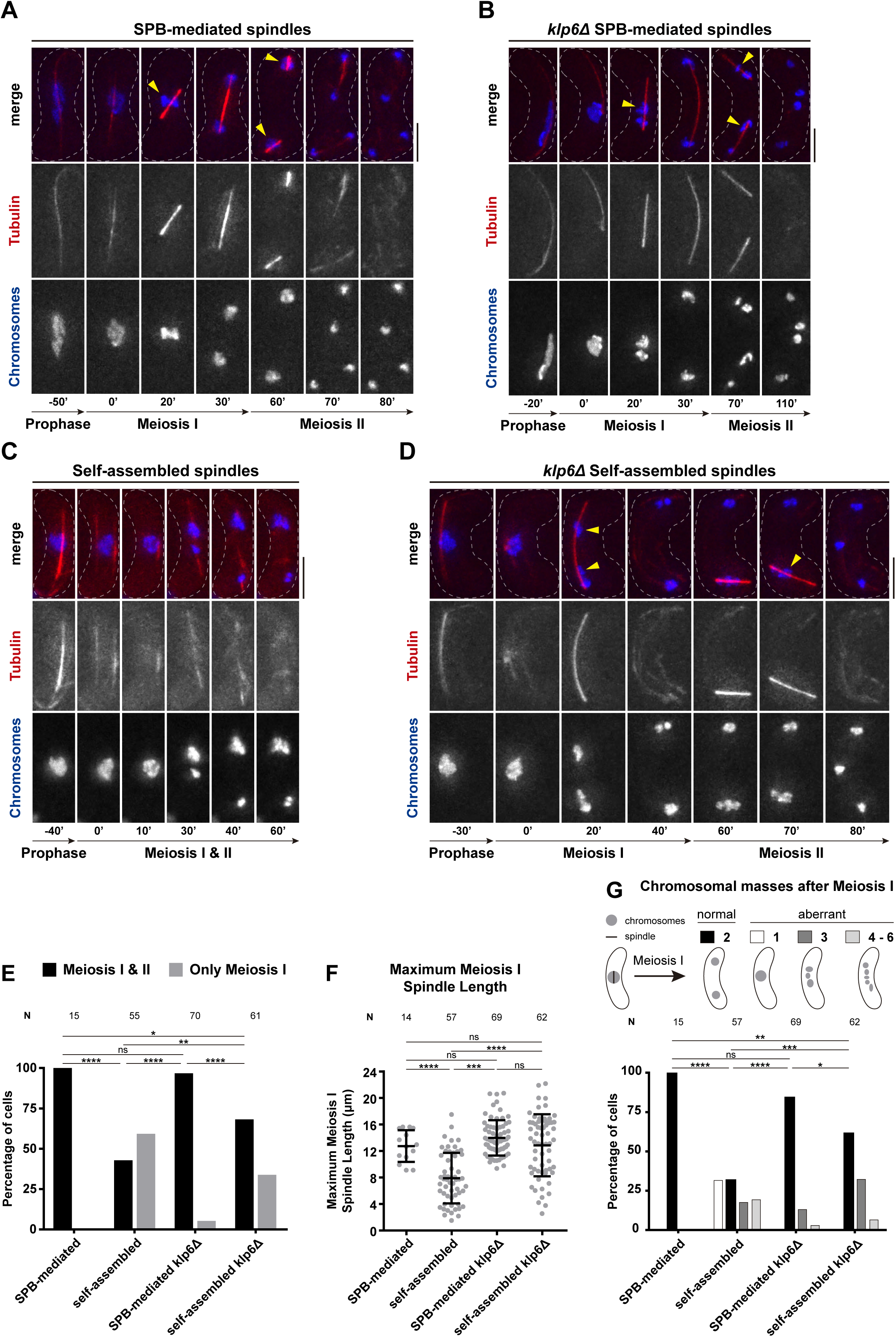
Kinesin-8 Klp6 deletion improves formation, structure and chromosome segregation of self-assembled spindles. **(A)** For SPB-mediated spindles, chromosomes position at the middle of the spindle body before being segregated equally to opposite spindle poles in MI (20’, yellow arrow) and MII (60’, yellow arrows). **(B)** In the absence of Klp6, chromosomes fail to congregate at the midzone of SPB-mediated spindles, instead remaining displaced towards one of the spindle poles for MI (20’, yellow arrowheads) and MII (70’, top yellow arrowhead). In addition, chromosome segregation defects, such as chromosome lagging are observed (70’, bottom arrowhead). **(C)** Self-assembled spindles present a thinner, more tenuous spindle body compared to SPB-mediated spindles (compare A 20’ vs C 30’) and segregate chromosomes deficiently (30’ to 60’). **(D)** Upon Klp6 deletion, self-assembled spindles present a thicker spindle body (compare C 30’ and D 20’), although chromosome mispositioning and lagging defects persist (20’ and 70’, yellow arrowheads). **(E)** Deletion of Klp6 increases the percentage of cells showing MII self-assembled spindles. Fisher’s exact test, *: p = 0.0161; **: p = 0.0015; ****: p < 0.0001. **(F)** Deletion of Klp6 enlarges MI self-assembled spindles maximum length up to a level indistinguishable from that of SPB-mediated spindles. t-test, ****: p < 0.0001; Mann-Whitney test, ***: p < 0.0001. **(G)** Deletion of Klp6 improves chromosome segregation fidelity by self-assembled spindles, rising the percentage of cells showing a number of chromosomal masses corresponding to normal segregation after meiosis I. Fisher’s exact test, *: p = 0.0052; **: p = 0.0036; ***: p = 0.0017.

Next, we analysed the rate of chromosome segregation defects in self-assembled spindles in cells with and without Klp6. In particular, we quantified the number of chromosome masses resulting after MI segregation. Most of *klp6Δ* SPB-dependent spindles segregate parental chromosomes into two masses (84%), although a minority exhibits three to six masses, consistent with chromosome segregation defects intrinsic to the loss of Klp5/6 (Fig. 7 G, n=69) (Pinder et al., 2019; Syrovatkina et al., 2013; Garcia et al., 2002; West et al., 2001). Self-assembled spindles present severe segregation defects: nearly a third (32%) of the spindles do not segregate chromosomes, leaving the parental nucleus as a single mass, while another third (36%) segregate chromosomes up to three to six masses and only a minority (32%) segregates into two masses i.e., normal segregation (Fig. 7 G, n=57). Strikingly, after *klp6* deletion, the rate of chromosome missegregation decreases, from 32% to 0% for one mass and from 19% to 6% for four to six masses. Consequently, the percentage of normal segregation rises from 32% to 61% as well as that of mild missegregation, i.e. three masses, from 18% to 32% (Fig. 7 G). Hence, *klp6* deletion increases chromosome segregation fidelity of self-assembled spindles, illustrating that spindle improvement occurs not only at the level of formation and structure, but also at the functional level.

## Discussion

In this work, we carried out a molecular characterization of an unexpected type of self-assembled spindle in fission yeast meiosis. SPB-dependent and independent spindles share similarities such as structural dependence on microtubule crosslinker Ase1/PRC1, microtubule arrangement polarity and the recruitment of the microtubule polymerase Alp14/XMAP215. In contrast, self-assembled spindle formation seems to be independent of the LINC complex localization and γ-tubulin complex conventional nucleation. Moreover, we increased self-assembled spindles robustness by deleting kinesin-8 *klp6*, which improved their structural stability and chromosome segregation fidelity. Thus, we think that our system will be useful for future studies to understand the molecular basis of the acentrosomal meiosis.

### Self-assembly of SPB-independent microtubules behaves as a proper spindle

A major question in our work is whether the self-assembly of nuclear microtubules in the absence of SPB insertion into the NE follows the characteristics of SPB-mediated spindles or, alternatively, are stochastically organised differing of the properties of the canonical spindle formation. Here, we confirm the presence of spindle-like properties; for instance, we have shown that self-assembled spindles require the canonical microtubule crosslinker Ase1/PRC1 to maintain their structural stability. Ase1 regulates the dynamics of both interphase and mitotic/meiotic microtubule arrays; nonetheless, deletion of Ase1 causes misorientation and misconfiguration of interphase arrays (Staub et al., 2005) while, in contrast, its deletion in mitosis leads to breakage of the spindle body at late stages of spindle elongation (Yamashita et al., 2005), consistently to what we observed for both meiotic SPB-mediated and self-assembled spindles. Congruently with our data, recent observations have well-established that Ase1 and microtubule motors together are enough to form spindle-like structures independently of the presence of MTOCs (Hannabuss et al., 2019; Blackwell et al., 2017).

Among the wide range of configurations permitted by microtubules versatility, the structure and polarity adopted by self-assembled microtubules repeats the scheme of SPB-dependent spindles, with microtubules minus-end at the spindle poles as shown by the recapitulating localization pattern of Pkl1. We hypothesize that self-assembled spindle polarity is the same than centrosomal spindles due to a set of microtubule-associated factors that force microtubules to adopt the canonical spindle-type configuration instead of alternative configurations, such as that of interphase arrays. Microtubules configuration within the spindle has fundamental implications for the mechanism of chromosome segregation. With minus-ends at the poles facing outwards and plus-ends projecting inwards to chromosomes, we speculate that the mechanism is likely to involve chromosome capturing by microtubules plus-ends and pulling to opposite poles driven by their shrinkage towards opposite poles, as for SPB-mediated spindles (Tanaka et al., 2005). In addition, the poleward movement of chromosomes that we observe could be complemented by supporting mechanisms such as pushing forces powered by microtubule polymerization (Ault et al., 1991), as reported for *C*. *elegans* meiosis (Laband et al., 2017), as well as microtubule sliding (Vukušić et al., 2017).

### Independence of self-assembled spindle formation from F-actin

In mammalian oocytes, F-actin cooperates with spindle microtubules to assist in polarization and maintenance of spindle structure (Roeles and Tsiavaliaris, 2019), drives the necessary asymmetrical spindle positioning within the oocyte (Mogessie et al., 2018) and helps ensure a correct chromosome segregation (Mogessie and Schuh, 2017). Given the importance of F-actin for mammalian oocytes, we decided to explore its relevance in the system of acentrosomal meiosis in fission yeast. Disruption of F-actin network in fission yeast meiosis uncovered that it is not required for self-assembled spindle formation nor dynamics.

Analysis of F-actin network during meiosis unveiled that self-assembled spindle formation shows an intrinsic defect in actin ring formation. In *S*. *pombe* meiosis, the SPBs localise at opposite poles of each MII nuclei and serve as reference point where the forespore membrane starts to form (Hirata and Shimoda, 1994; Hagan and Yanagida, 1995; Tanaka and Hirata, 1982); then, actin rings assemble and contract coordinately with MII nuclear divisions, guiding the extension and wrapping of the membrane around each nuclei (Yan and Balasubramanian, 2012; Itadani et al., 2006; Ohtaka et al., 2007). Given that the SPBs seem to trigger forespore membrane formation and in *bqt1Δ sad1*.*2* cells the SPBs are completely detached from the NE, actin rings may be unable to assemble properly around nuclei.

### How are the self-assembled spindles nucleated in fission yeast meiosis?

Our observations suggest that conventional γ-tubulin complex-driven nucleation may not be involved in self-assembled spindle formation since we did not observe accumulation of Alp4, one of the essential members of the complex at the self-assembled spindle structure. However, we cannot totally rule out this possibility. For instance, the accumulation of Alp4 molecules at the SPBs in a SPB-dependent spindle context might be necessary for triggering SPB insertion into the NE instead for the nucleation of the SPB-dependent spindles. Accordingly, Alp4-GFP dots show similar behaviour between inserted and uninserted SPBs (Fig. 5B, 75’, yellow asterisk), which strongly suggests that these dots correspond to Alp4 molecules associated to the SPBs independently of their association to the NE, and then, previous to the insertion process.

On the other hand, other alternative explanation to the absence of Alp4 signal in the vicinity of forming self-assembled spindle might derive from the fact that self-assembled spindles show a thinner structure than SPB-dependent spindles, which indicates a lower number of microtubules. We hypothesize that self-assembled spindles in our system would then need a smaller amount of nucleation factors, among them the γ-tubulin complex, than the canonical spindle. Hence, these factors could not be easily detectable by live fluorescence microscopy systems used in this study.

Together with nucleation of microtubules, another essential feature for spindle assembly is microtubule polymerization by polymerases, aiding SPB separation and spindle elongation (Yukawa et al., 2017), such as Alp14/XMAP215. The fact that Alp14 accumulates into the nucleus in self-assembled spindles formation in a similar manner than SPB-dependent spindles unveils that nucleocytoplasmic traffic of Alp14 in the absence of SPB insertion is normal. We think that this observation of proper transport can be generalized to other spindle-related factors. In particular, given the near-wildtype Alp14 behaviour and its functional dependence on its partner Alp7 (Sato et al., 2004), this latter may follow a wild-type-like behaviour as well, possibly localising to self-assembled spindle microtubules. In general, based on their nuclear localization observed in previous work (Pineda-Santaella and Fernández-Álvarez, 2019) and this study, this trend could also apply to Ase1, Pkl1 and components of the γ-tubulin complex. Hence, we believe that the feasibility of self-assembled spindle formation is firstly founded on the availability of such factors within the nuclear environment.

Another interesting feature of Alp14 behaviour is its rather dispersed distribution inside the nucleus at the time of self-assembled spindle formation compared to the SPB-mediated spindle setting. We propose that because self-assembled spindle might be composed of a fewer number of microtubules, fewer molecules of Alp14 would be loaded into the spindle body and thus, the rest of Alp14 molecules would stay free at the nucleoplasm unable to load into the saturated microtubule lattice. Considering Alp14 normal nuclear accumulation onset as a temporal reference, this uncovered a delay in self-assembled spindle formation compared with SPB-mediated spindles. We predict that spindle self-assembly mechanism is less efficient upon the lack of reference and organization from the SPBs and/or present higher requirements for spindle assembly factors, bearing an inherent defectiveness that translates into more time needed to form. Unfortunately, because *bqt1*Δ *sad1*.*2 alp14*Δ cells show growth polarity defects, typical of *alp* genes misfunction (Radcliffe et al., 1998) and severe growth defects, it is difficult to explore the consequences of the absence of Alp14 in self-assembled spindles during fission yeast meiosis. We think that this growth impairment is consequence of the combination of *alp14* and *sad1*.*2* misfunctions, which is in concordance to the described genetic interaction between *alp14* and *csi1*, a partner of Sad1, necessary to stabilize centromere/telomere-Sad1 interactions and the correct spindle formation and chromosome segregation (Hou et al., 2012).

### Why Kinesin-8 deletion improves self-assembled spindles?

Elimination of the kinesin-8 Klp6 increases self-assembled spindle microtubules stability, and consequently, efficiency of chromosome segregation in *bqt1Δ sad1*.*2* cells. One of the roles of Kinesin-8 Klp5/Klp6 is to destabilize microtubules by i) promoting microtubule catastrophe (Erent et al., 2012; Unsworth et al., 2008) and ii) hamper polymerization and enhance depolymerization (Garcia et al., 2002). A possible mechanism for the increase in thickness of self-assembled spindles is enhanced polymerization and elongation of tubulin subunits incorporated into the lattice of already formed microtubules filaments (Schaedel et al., 2019), resulting in new filaments which widen the spindle body (Fig. 8 A and B). Longer length of *klp6*-deleted self-assembled spindles could be explained by microtubule overgrowth derived from enhanced polymerization, as well as by microtubule hyper-stabilization, which would confer the spindle resistance against microtubule-destabilizing mechanisms responsible for spindle disassembly, letting it reach a longer length (Fig. 8 A and B). Enhanced polymerization would as well promote self-assembled spindle formation, explaining the increase in the appearance of self-assembled spindles in MII (Fig. 8 B and C). Regarding chromosome segregation, loss of Klp6 leads to chromosome segregation defects in a SPB-mediated setting (Pinder et al., 2019; Syrovatkina et al., 2013; Garcia et al., 2002; West et al., 2002) (Fig. 7). However, in our system, deletion of Klp6 significantly improves chromosome segregation carried out by self-assembled spindles, probably as a consequence of reinforced spindle structure and kinetochore-microtubule interaction stability, which might facilitate kinetochore attachment (Fig. 8 B). Hence, our results show a successful strategy aimed to improve the acentrosomal spindle formation and function (Fig. 8 C), which we estimate not only relevant for future studies in fission yeast meiosis but also to gain insight into mammalian female meiosis.

**FIGURE 8.**
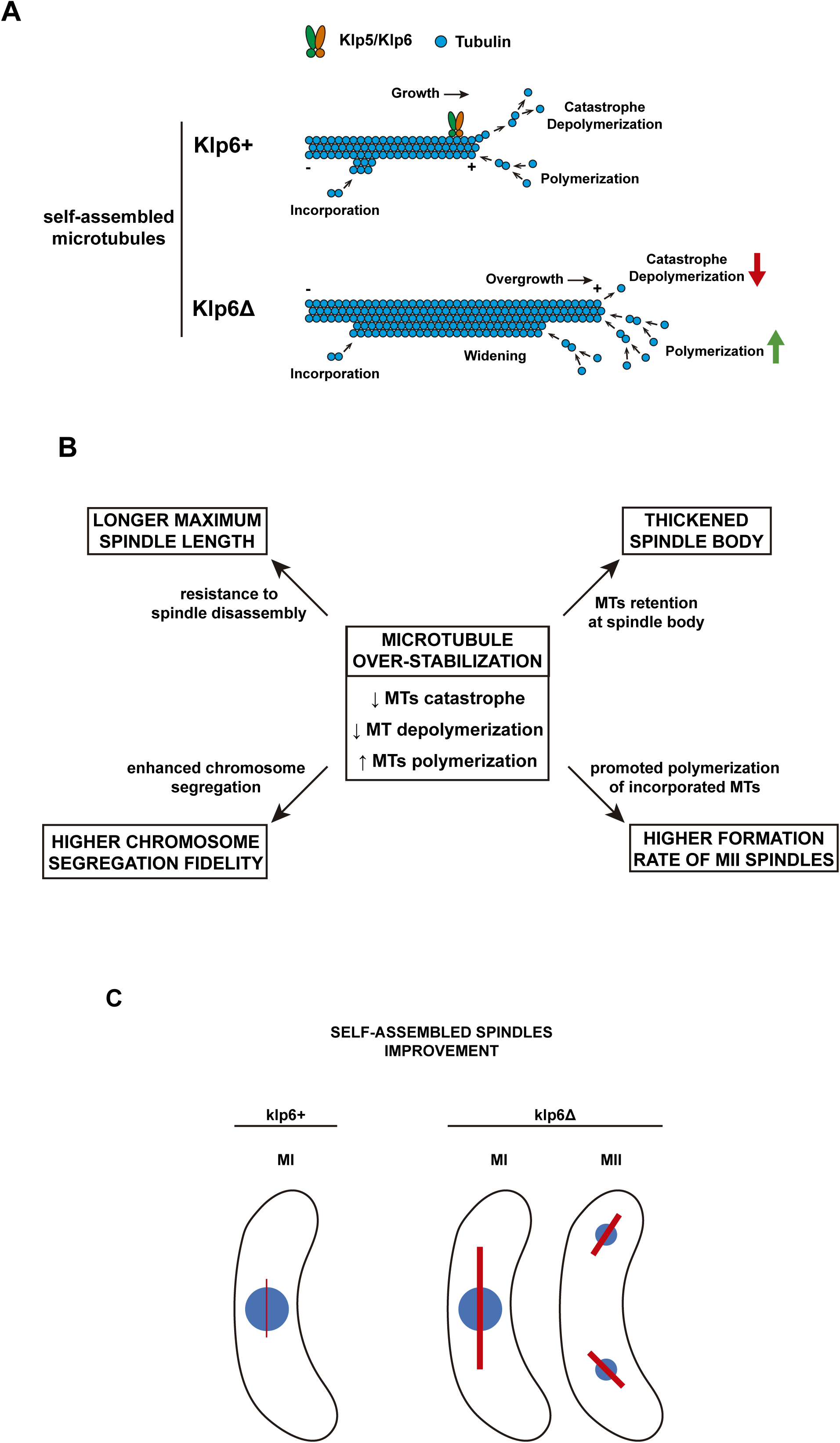
Possible mechanisms of self-assembled spindle improvement by Klp6 deletion. **(A)** Elimination of microtubule destabilizing activity of Klp5/Klp6 (*klp6Δ*) shifts microtubule dynamics equilibrium towards increased polymerization and decreased catastrophe and depolymerisation. This translates into enhanced polymerization from microtubule filament plus-ends, leading to microtubule overgrowth, as well as from free tubulin incorporated into the microtubule lattice, leading to microtubule widening. **(B)** Microtubule over-stabilization caused by Klp6 deletion may derive into secondary effects responsible for self-assembled spindles improvement through different pathways. **(C)** Macroscopic effect of self-assembled spindle improvement consists of a reinforcement of spindle structure, promotion of MII spindle formation and increase of chromosome segregation fidelity.

## Materials and methods

### Strains culture and meiosis induction

Strains used in this work are listed in Supplementary Table 1. Strains were thawed in solid YES rich medium plate and incubated for 24 h at 32°C or 48 h at 25°C and patched in a new fresh plate and incubated for 24 h at 32°C or 25°C. For meiosis induction, strains were repatched in solid sporulating SPA medium and incubated for 6-6.5 h at 28°C, the optimal temperature for sporulation. After induction, meiotic cells are immobilised with lectin (0.2 µg/mL, Sigma Aldrich, L1395) at the bottom of a µ-Dish glass-bottom 35 mm uncoated dish (Cat. No: 81151, ibidi Gmbh), washed and covered in a total of 3 mL of EMM2 minimal medium without nitrogen with or without drug to assure continuation of meiosis.

### Live microscopy images acquisition and processing

For acquisition of live fluorescence microscopy images, two microscopy systems were used: Spinning Disk Confocal Microscope (Photometrics Evolve camera; Olympus 100x 1.4 NA oil immersion objective; Roper Scientific-Photometrics) and Delta Vision (CoolSnap HQ camera; Olympus 100x 1.4 NA oil immersion objective, Environmental temperature and CO_2_ precision control; Inverted Microscope Olympus IX71). For Spinning Disk, images were taken using 14 z-sections separated by 0.5 µm during 5-6 hours every 5 minutes, with the following channels, exposure times and laser intensities: mCherry (561 nm), 150 ms, 50%; GFP (491 nm), 100 ms, 25%; CFP (405 nm), 100 ms, 20%; Brightfield (visible), 50 ms. For Delta Vision, images were taken using 15-20 z-sections separated by 0.4 µm during 5-6 hours every 5 or 10 minutes, with the following channels, exposure times and radiation intensities: YFP (492 nm), 150ms, 50%; CFP (436 nm), 100 ms, 32%; TRITC (555 nm), 500 ms, 100%; Brightfield (visible), 200 ms, 10%. Maximum Z-projections of acquired images were obtained and stacked with ImageJ software (version 1.52p) (http://rsbweb.nih.gov/ij/). Acquired images were deconvolved with the PSFs and scripts available at https://github.com/danilexn/deCU, based on the Richardson-Lucy algorithm implementation by Shao and Milkie, at Betzig lab (https://github.com/scopetools/cudaDecon). Further image processing was performed using Adobe Photoshop CC 2018 (Adobe Inc.) and Adobe Illustrator CC 2017 (Adobe Inc.). Only meiotic cells which progressed normally into and along meiosis were submitted to analysis, discarding cells with (pre-)meiotic defects, such as non-fusion of parental nuclei (karyogamy), or viability defects, such as cell death during image acquisition.

### Latrunculin-A treatment

To treat meiotic cells with Latrunculin-A (Sigma-Aldrich, L5163), 10 µL of 1 mM Latrunculin-A were mixed with 490 µL of EMM2 minimal medium and these 500 µL were mixed with the remaining 2.5 mL of EMM2 minimal medium used to cover cells during image acquisition. This manner, the final concentration of Latrunculin-A in the total 3 mL of medium is 4 µM. Meiotic cells were added Latrunculin A after meiosis induction and incubated with the drug during the whole time of filming.

### Quantification and statistical analysis

Quantification of nuclear fluorescence signal of Alp14-GFP was performed by measuring in a maximum Z-projection the signal intensity within the nuclear environment, delimited by the perimeter described by the signal of the parental nucleus chromosomal mass before spindle formation.

Statistical tests were performed with GraphPad Prism 6. To test for difference between the mean of two distributions, if both followed a normal distribution, a parametric Student’s t-test was performed; otherwise, a non-parametric Mann-Whitney test was performed. To test for difference between two proportions, a Fisher’s exact test was performed.

## Supplemental material

## Acknowledgements

We thank Sergio Rincón and Ana Sánchez Molina for critical comments on the manuscript; Daniel León-Periñán for digital image processing; Alejandra Cano for technical support; and the CABD microscopy facility technician Katherina García. We would like to thank the Genetics Department and Springboard lab for their useful discussion and comments, especially Víctor Carranco for technical support. This work was supported by Spanish Government, Plan Nacional PGC2018-098118-A-I00 and Ramon y Cajal program, RyC-2016-19659 to AF-A; and by the Pablo de Olavide University “Ayuda Puente Predoctoral” fellowship (PPI1803) to AP-S. The CABD is an institution funded by Pablo de Olavide University, Consejo Superior de Investigaciones Científicas (CSIC) and Junta de Andalucía.

## Author contributions

AP-S. and AF-A. designed the study; AP-S. performed all the experiments with the support of NF-C. and AS-G.; AF-A. acquired funding and supervised the project; AP-S and AF-A. wrote the manuscript.

## Abbreviations

MI: meiosis I
MII: meiosis II
MTOC: microtubule-organizing centre
NE: nuclear envelope
SPB: spindle pole body

---

**FIGURE S1.**
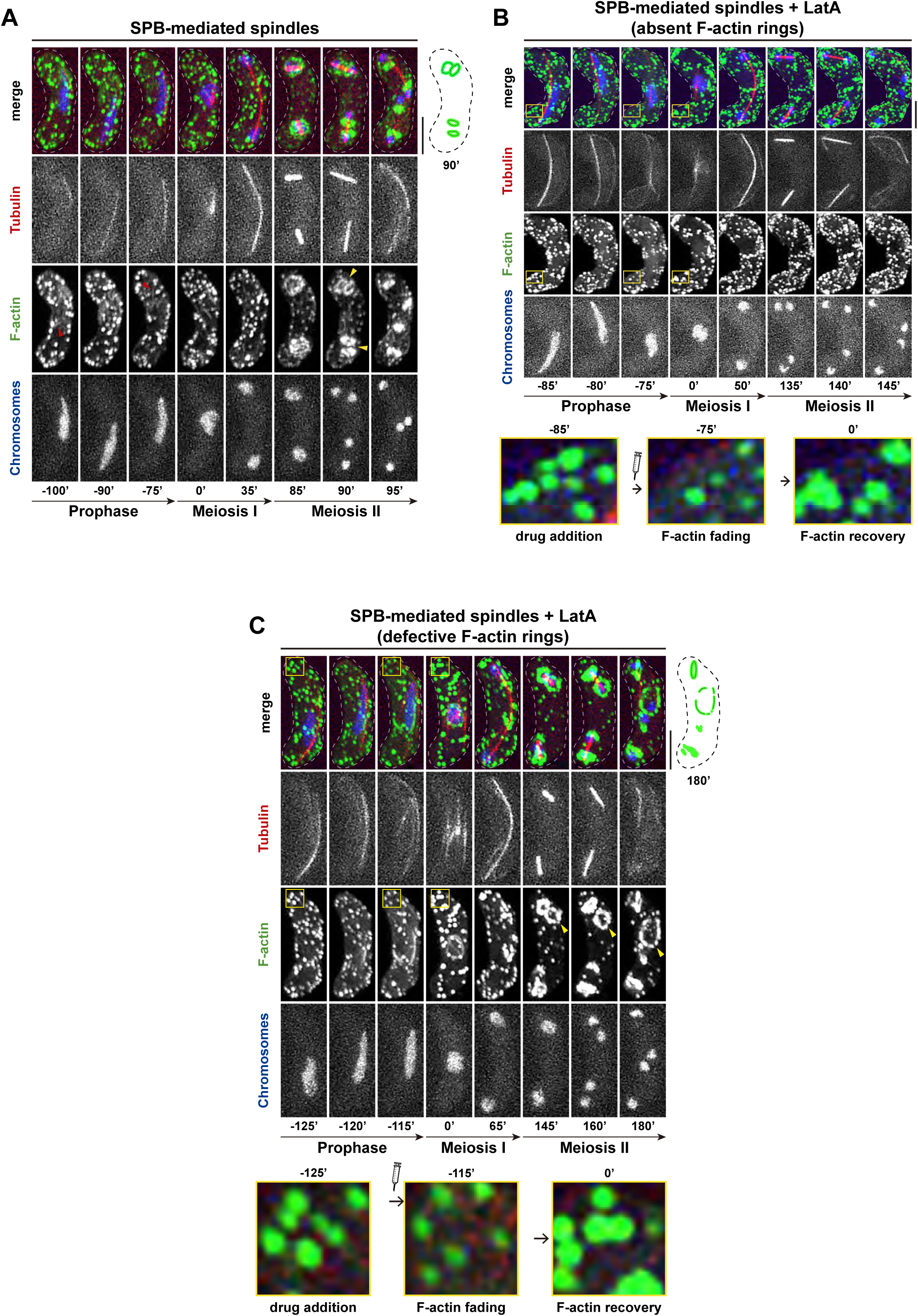
SPB-mediated meiotic spindle formation and dynamics are independent of the F-actin network in fission yeast meiosis. **(A)** In meiotic prophase, microtubules drag the nucleus in their oscillating movements, process called horsetail stage (−100’ to −75’). After prophase, MI (0’ to 35’) and MII (85’ to 95’) SPB-mediated spindles form. Unaltered F-actin network form dispersed patches and thin cables (red arrowheads) during prophase, while in MII it forms 2 rings around each daughter nuclei resulting from MI division (85’ to 95’). **(B, C)** Upon addition of actin-depolymerizing drug Latrunculin A (LatA, 4 µM), actin cables are no longer observable and F-actin patches signal is reduced and later recovered (zoomed areas indicated with rectangles). Actin rings either **(B)** do not form in MII (135’ to 145’) or **(C)** show aberrant morphology and dynamics (145’ to 180’, yellow arrowhead). In spite of F-actin disruption, MI and II spindles still form and display wildtype-like morphology and dynamics. F-actin: Lifeact-GFP. Cartoons: schematic representation of indicated frames.

| Strain | Mating Type | Genotype | Origin |
| --- | --- | --- | --- |
| AFA51 | *h<sup>90</sup>* | *ade6-M210 ura4-D18 hht1-CFP:his3 Pnda3-mCherry-Atb2:aur1* | This study |
| AFA54 | *h<sup>90</sup>* | *ade6-M210 ura4-D18 hht1-CFP:his3 Pnda3-mCherry-Atb2:aur1 sad1.T3S.S52P-13Myc-hphMX6 bqt1::hygMX6* | This study |
| AFA119 | *h<sup>90</sup>* | *ade6-M210 hht1-CFP:his3 Pnda3-mCherry-Atb2:aur1 Pact1-LA-GFP:leu1+* | This study |
| AFA210 | *h<sup>90</sup>* | *his3-D1 hht1-CFP:his3+ Pnda3-mCherry-atb2:aur1 alp4-GFP:kanMX6* | This study |
| AFA219 | *h<sup>90</sup>* | *hht1-CFP:his3+ Pnda3-mCherry-atb2:aur1 alp4-GFP:kanMX6 sad1.T3S.S52P-13Myc-hphMX6 bqt1::hygMX6* | This study |
| AFA306 | *h<sup>90</sup>* | *ade6-M210 ura4-D18 hht1-CFP:his3 Pnda3-mCherry-Atb2:aur1 ase1Δ::KanMX6* | This study |
| AFA316 | *h<sup>90</sup>* | *ade6-M210 ura4-D18 hht1-CFP:his3 Pnda3-mCherry-Atb2:aur1 sad1.T3S.S52P-13Myc-hphMX6 bqt1::hygMX6 ase1Δ::KanMX6* | This study |
| AFA319 | *h<sup>90</sup>* | *hht1-CFP:his3 Pnda3-mCherry-Atb2:aur1 sad1.T3S.S52P-13Myc-hphMX6 bqt1::hygMX6 Pact1-LA-GFP:leu1+* | This study |
| AFA336 | *h<sup>90</sup>* | *hht1-CFP:his3 Pnda3-mCherry-Atb2:aur1 alp14-GFP:Kan* | This study |
| AFA339 | *h<sup>90</sup>* | *his3-D1 hht1-CFP:his3 Pnda3-mCherry-Atb2:aur1 sad1.T3S.S52P-13Myc-hphMX6 bqt1::hygMX6 alp14-GFP:Kan* | This study |
| AFA350 | *h<sup>90</sup>* | *kanR-Palp4-GFP-pkl1 hht1-CFP:his3 aur1R-Pnda3-mCherry-atb2* | This study |
| AFA406 | *h<sup>90</sup>* | *kanR-Palp4-GFP-pkl1 hht1-CFP:his3 Pnda3-mCherry-Atb2:aur1 sad1.T3S.S52P-13Myc-hphMX6 bqt1::hygMX6* | This study |
| AFA416 | *h<sup>90</sup>* | *ade6-M210 hht1-CFP:his3+ Pnda3-mCherry-Atb2:aur1 sad1-GFP:HygMX6* | This study |
| AFA437 | *h<sup>90</sup>* | *ade6-M210 hht1-CFP:his3+ Pnda3-mCherry-atb2:aur1 bqt1::hygMX6 sad1.2-GFP:KanMX6* | This study |
| AFA438 | *h<sup>90</sup>* | *hht1-CFP:his3+ Pnda3-mCherry-Atb2:aur1 sad1.2-GFP:HygMX6* | This study |
| AFA1431 | *h<sup>90</sup>* | *ade6- hht1-CFP:his3 Pnda3-mCherry-Atb2:aur1 klp6::kanMX6* | This study |
| AFA1437 | *h<sup>90</sup>* | *hht1-CFP:his3 Pnda3-mCherry-Atb2:aur1 sad1.T3S.S52P::natMX6 bqt1::hygMX6 klp6::kanMX6* | This study |
| JCF10327 | *h<sup>90</sup>* | *alp4-GFP-kanMX6 sid4-mRFP:hygMX6 hrs1::natMX6* | Lab stock |
| MY858 | *h<sup>-</sup>* | *kanR-Palp4-GFP-pkl1 aur1R-Pnda3-mCherry-atb2 leu1 ura4* | Yukawa et al., 2019 |
| NF771 | *h<sup>+</sup>* | *alp14-GFP:Kan* | Lab stock |
| RD234 | *h<sup>+</sup>* | *ase1Δ::KanMX6* | Lab stock |
| RD5948 | *h<sup>-</sup>* | *ura4-D18 leu1-32 Pact1-LA-GFP:leu1+* | Lab stock |
| SR1.33 | *h<sup>+</sup>* | *klp6::kanMX6 leu1-32 ura4-D18 ade6-* | Lab stock |

## References

Al-Bassam, J., H. Kim, I. Flor-Parra, N. Lal, H. Velji, and F. Chang. 2012. Fission yeast Alp14 is a dose-dependent plus end-tracking microtubule polymerase. Mol. Biol. Cell. 23:2878–2890. doi: 10.1091/mbc.E12-03-0205.

Ault, J.G., A.J. Demarco, E.D. Salmon, and C.L. Rieder. 1991. Studies on the ejection properties of asters: Astral microtubule turnover influences the oscillatory behavior and positioning of mono-oriented chromosomes. J. Cell Sci. 99:701–710.

Bestul, A.J., Z. Yu, J.R. Unruh, and S.L. Jaspersen. 2017. Molecular model of fission yeast centrosome assembly determined by superresolution imaging. J. Cell Biol. 216:2409–2424. doi: 10.1083/jcb.201701041.

Bettencourt-Dias, M., and D.M. Glover. 2007. Centrosome biogenesis and function: Centrosomics brings new understanding. Nat. Rev. Mol. Cell Biol. 8:451–463. doi: 10.1038/nrm2180.

Blackwell, R., C. Edelmaier, O. Sweezy-Schindler, A. Lamson, Z.R. Gergely, E. O’Toole, A. Crapo, L.E. Hough, J.R. McIntosh, M.A. Glaser, and M.D. Betterton. 2017. Physical determinants of bipolar mitotic spindle assembly and stability in fission yeast. Sci. Adv. 3. doi: 10.1126/sciadv.1601603.

Bratman, S. V., and F. Chang. 2007. Stabilization of Overlapping Microtubules by Fission Yeast CLASP. Dev. Cell. 13:812–827. doi: 10.1016/j.devcel.2007.10.015.

Braunfeld, M.B., M. Moritz, D.A. Agard, J. Heuser, and V. Guénebaut. 2002. Structure of the γ-tubulin ring complex: a template for microtubule nucleation. Nat. Cell Biol. 2:365–370. doi: 10.1038/35014058.

Chikashige, Y., C. Tsutsumi, M. Yamane, K. Okamasa, T. Haraguchi, and Y. Hiraoka. 2006. Meiotic Proteins Bqt1 and Bqt2 Tether Telomeres to Form the Bouquet Arrangement of Chromosomes. Cell. 125:59–69. doi: 10.1016/j.cell.2006.01.048.

Clift, D., and M. Schuh. 2015. A three-step MTOC fragmentation mechanism facilitates bipolar spindle assembly in mouse oocytes. Nat. Commun. 6:1–12. doi: 10.1038/ncomms8217.

Colombié, N., C.F. Cullen, A.L. Brittle, J.K. Jang, W.C. Earnshaw, M. Carmena, K. McKim, and H. Ohkura. 2008. Dual roles of incenp crucial to the assembly of the acentrosomal metaphase spindle in female meiosis. Development. 135:3239–3246. doi: 10.1242/dev.022624.

Cooley, L., and W.E. Theurkauf. 1994. Cytoskeletal functions during Drosophila oogenesis. Science (80-.). 266:590–596. doi: 10.1126/science.7939713.

Ding, R., R.R. West, M. Morphew, B.R. Oakley, and J. Richard McIntosh. 1997. The spindle pole body of Schizosaccharomyces pombe enters and leaves the nuclear envelope as the cell cycle proceeds. Mol. Biol. Cell. 8:1461–1479. doi: 10.1091/mbc.8.8.1461.

Dumont, J., and A. Desai. 2012. Acentrosomal spindle assembly and chromosome segregation during oocyte meiosis. Trends Cell Biol. 22:241–249. doi: 10.1016/j.tcb.2012.02.007.

Ebina, H., L. Ji, and M. Sato. 2019. CLASP promotes microtubule bundling in metaphase spindle independently of Ase1/PRC1 in fission yeast. Biol. Open. 8. doi: 10.1242/bio.045716.

Erent, M., D.R. Drummond, and R.A. Cross. 2012. S. pombe kinesins-8 promote both nucleation and catastrophe of microtubules. PLoS One. 7. doi: 10.1371/journal.pone.0030738.

Fennell, A., A. Fernández-álvarez, K. Tomita, and J.P. Cooper. 2015. Telomeres and centromeres have interchangeable roles in promoting meiotic spindle formation. J. Cell Biol. 208:415–428. doi: 10.1083/jcb.201409058.

Fernández-Álvarez, A., C. Bez, E.T. O’Toole, M. Morphew, and J.P. Cooper. 2016. Mitotic Nuclear Envelope Breakdown and Spindle Nucleation Are Controlled by Interphase Contacts between Centromeres and the Nuclear Envelope. Dev. Cell. 39:544–559. doi: 10.1016/j.devcel.2016.10.021.

Flor-Parra, I., A.B. Iglesias-Romero, and F. Chang. 2018. The XMAP215 Ortholog Alp14 Promotes Microtubule Nucleation in Fission Yeast. Curr. Biol. 28:1681–1691.e4. doi: 10.1016/j.cub.2018.04.008.

Gachet, Y., C. Reyes, S. Goldstone, and S. Tournier. 2006. The fission yeast spindle orientation checkpoint: a model that generates tension? Yeast. 23:1015–1029. doi: 10.1002/yea.1410.

Gachet, Y., S. Tournier, J.B.A. Millar, and J.S. Hyams. 2001. A MAP kinase-dependent actin checkpoint ensures proper spindle orientation in fission yeast. Nature. 412:352–355. doi: 10.1038/35085604.

Garcia, M.A. 2001. Fission yeast ch-TOG/XMAP215 homologue Alp14 connects mitotic spindles with the kinetochore and is a component of the Mad2-dependent spindle checkpoint. EMBO J. 20:3389–3401. doi: 10.1093/emboj/20.13.3389.

Garcia, M.A., N. Koonrugsa, and T. Toda. 2002. Spindle-kinetochore attachment requires the combined action of Kin I-like Klp5/6 and Alp14/Dis1-MAPs in fission yeast. EMBO J. 21:6015–6024. doi: 10.1093/emboj/cdf611.

Gard, D.L. 1991. Organization, nucleation, and acetylation of microtubules in Xenopus laevis oocytes: A study by confocal immunofluorescence microscopy. Dev. Biol. 143:346–362. doi: 10.1016/0012-1606(91)90085-H.

Gergely, Z.R., A. Crapo, L.E. Hough, J. Richard McIntosh, and M.D. Betterton. 2016. Kinesin-8 effects on mitotic microtubule dynamics contribute to spindle function in fission yeast. Mol. Biol. Cell. 27:3490–3514. doi: 10.1091/mbc.E15-07-0505.

Gonzalez, C., G. Tavosanis, and C. Mollinari. 1998. Centrosomes and microtubule organisation during Drosophila development. J. Cell Sci. 111:2697–2706.

Grallert, A., C. Beuter, R.A. Craven, S. Bagley, D. Wilks, U. Fleig, and I.M. Hagan. 2006. S. pombe CLASP needs dynein, not EB1 or CLIP170, to induce microtubule instability and slows polymerization rates at cell tips in a dynein-dependent manner. Genes Dev. 20:2421–2436. doi: 10.1101/gad.381306.

Griswold, M.D., and P.A. Hunt. 2013. Meiosis. In Brenner’s Encyclopedia of Genetics: Second Edition.

Hagan, I., and M. Yanagida. 1995. The product of the spindle formation gene sad1+ associates with the fission yeast spindle pole body and is essential for viability. J. Cell Biol. 129:1033–1047. doi: 10.1083/jcb.129.4.1033.

Hannabuss, J., M. Lera-Ramirez, N.I. Cade, F.J. Fourniol, F. Nédélec, and T. Surrey. 2019. Self-Organization of Minimal Anaphase Spindle Midzone Bundles. Curr. Biol. 29:2120–2130.e7. doi: 10.1016/j.cub.2019.05.049.

Heald, R., R. Tournebize, T. Blank, R. Sandaltzopoulos, P. Becker, A. Hyman, and E. Karsenti. 1996. Self-organization of microtubules into bipolar spindles around artificial chromosomes in Xenopus egg extracts. Nature. 382:420–425. doi: 10.1038/382420a0.

Hertig, A.T., and E.C. Adams. 1967. STUDIES ON THE HUMAN OOCYTE AND ITS FOLLICLE. J. Cell Biol. 34:647–675. doi: 10.1083/jcb.34.2.647.

Hirata, A., and C. Shimoda. 1994. Structural modification of spindle pole bodies during meiosis II is essential for the normal formation of ascospores in Schizosaccharomyces pombe: ultrastructural analysis of spo mutants. Yeast. 10:173–183. doi: 10.1002/yea.320100205.

Holubcova, Z., M. Blayney, K. Elder, and M. Schuh. 2015. Error-prone chromosome-mediated spindle assembly favors chromosome segregation defects in human oocytes. Science (80-.). 348:1143–1147. doi: 10.1126/science.aaa9529.

Horio, T., S. Uzawa, M.K. Jung, B.R. Oakley, K. Tanaka, and M. Yanagida. 1991. The fission yeast γ-tubulin is essential for mitosis and is localized at microtubule organizing centres. J. Cell Sci. 99:693–700.

Hou, H., Z. Zhou, Y. Wang, J. Wang, S.P. Kallgren, T. Kurchuk, E.A. Miller, F. Chang, and S. Jia. 2012. Csi1 links centromeres to the nuclear envelope for centromere clustering. J. Cell Biol. 199:735–744. doi: 10.1083/jcb.201208001.

Itadani, A., T. Nakamura, and C. Shimoda. 2006. Localization of Type I Myosin and F-actin to the Leading Edge Region of the Forespore Membrane in Schizosaccharomyces pombe. 195:181–195.

Kemp, C.A., K.R. Kopish, P. Zipperlen, J. Ahringer, and K.F. O’Connell. 2004. Centrosome maturation and duplication in C. elegans require the coiled-coil protein SPD-2. Dev. Cell. 6:511–523. doi: 10.1016/S1534-5807(04)00066-8.

Kolano, A., S. Brunet, A.D. Silk, D.W. Cleveland, and M.H. Verlhac. 2012. Error-prone mammalian female meiosis from silencing the spindle assembly checkpoint without normal interkinetochore tension. Proc. Natl. Acad. Sci. U. S. A. 109. doi: 10.1073/pnas.1204686109.

Laband, K., R. Le Borgne, F. Edwards, M. Stefanutti, J.C. Canman, J.M. Verbavatz, and J. Dumont. 2017. Chromosome segregation occurs by microtubule pushing in oocytes. Nat. Commun. 8:1–10. doi: 10.1038/s41467-017-01539-8.

Longo, F.J., and E. Anderson. 1969. Cytological aspects of fertilization in the lamellibranch, Mytilus edulis Polar body formation and development of the female pronucleus. J. Exp. Zool. 172:69–95. doi: 10.1002/jez.1401720107.

Masuda, H., R. Mori, M. Yukawa, and T. Toda. 2013. Fission yeast MOZART1/Mzt1 is an essential γ-tubulin complex component required for complex recruitment to the microtubule organizing center, but not its assembly. Mol. Biol. Cell. 24:2894–2906. doi: 10.1091/mbc.E13-05-0235.

Mogessie, B., K. Scheffler, and M. Schuh. 2018. Assembly and Positioning of the Oocyte Meiotic Spindle. Annu. Rev. Cell Dev. Biol. 34:381–403. doi: 10.1146/annurev-cellbio-100616-060553.

Mogessie, B., and M. Schuh. 2017. Actin protects mammalian eggs against chromosome segregation errors. Science (80-.). 357. doi: 10.1126/science.aal1647.

Nakashima, S., and K.H. Kato. 2001. Centriole behavior during meiosis in oocytes of the sea urchin Hemicentrotus pulcherrimus. Dev. Growth Differ. 43:437–445. doi: 10.1046/j.1440-169X.2001.00580.x.

Niwa, O., M. Shimanuki, and F. Miki. 2000. Telomere-led bouquet formation facilitates homologous chromosome pairing and restricts ectopic interaction in fission yeast meiosis. EMBO J. 19:3831–3840. doi: 10.1093/emboj/19.14.3831.

Ohtaka, A., D. Okuzaki, T.T. Saito, and H. Nojima. 2007. Mcp4, a meiotic coiled-coil protein, plays a role in F-actin positioning during Schizosaccharomyces pombe meiosis. Eukaryot. Cell. 6:971–983. doi: 10.1128/EC.00016-07.

Pidoux, A.L., M. LeDizet, and W.Z. Cande. 1996. Fission yeast pkl1 is a kinesin-related protein involved in mitotic spindle function. Mol. Biol. Cell. 7:1639–1655. doi: 10.1091/mbc.7.10.1639.

Pinder, C., Y. Matsuo, S.P. Maurer, and T. Toda. 2019. Kinesin-8 and Dis1/TOG collaborate to limit spindle elongation from prophase to anaphase A for proper chromosome segregation in fission yeast. J. Cell Sci. 132. doi: 10.1242/jcs.232306.

Pineda-Santaella, A., and A. Fernández-Álvarez. 2019. Spindle assembly without spindle pole body insertion into the nuclear envelope in fission yeast meiosis. Chromosoma. 128:267–277. doi: 10.1007/s00412-019-00710-y.

Radcliffe, P., D. Hirata, D. Childs, L. Vardy, and T. Toda. 1998. Identification of novel temperature-sensitive lethal alleles in essential β-tubulin and nonessential α2-tubulin genes as fission yeast polarity mutants. Mol. Biol. Cell. 9:1757–1771. doi: 10.1091/mbc.9.7.1757.

Riedl, J., A.H. Crevenna, K. Kessenbrock, J.H. Yu, D. Neukirchen, M. Bista, F. Bradke, D. Jenne, T.A. Holak, Z. Werb, M. Sixt, and R. Wedlich-Soldner. 2008. Lifeact: A versatile marker to visualize F-actin. Nat. Methods. 5:605–607. doi: 10.1038/nmeth.1220.

Roeles, J., and G. Tsiavaliaris. 2019. Actin-microtubule interplay coordinates spindle assembly in human oocytes. Nat. Commun. 10:1–10. doi: 10.1038/s41467-019-12674-9.

Rothballer, A., T.U. Schwartz, and U. Kutay. 2013. LINCing complex functions at the nuclear envelope what the molecular architecture of the LINC complex can reveal about its function. Nucl. (United States). 4:1–8. doi: 10.4161/nucl.23387.

Ruthmann, A. 1959. The Fine Structure of the Meiotic Spindle of the Crayfish. J. Biophys. Biochem. Cytol. 5:177–179. doi: 10.1083/jcb.5.1.177.

Sathananthan, A.H., K. Selvaraj, M. Lakshmi Girijashankar, V. Ganesh, P. Selvaraj, and A.O. Trounson. 2006. From oogonia to mature oocytes: Inactivation of the maternal centrosome in humans. Microsc. Res. Tech. 69:396–407. doi: 10.1002/jemt.20299.

Sato, M., L. Vardy, M.A. Garcia, N. Koonrugsa, and T. Toda. 2004. Interdependency of Fission Yeast Alp14 / TOG and Coiled Coil Protein Alp7 in Microtubule Localization and Bipolar Spindle Formation □. 15:1609–1622. doi: 10.1091/mbc.E03.

Schaedel, L., S. Triclin, D. Chrétien, A. Abrieu, C. Aumeier, J. Gaillard, L. Blanchoin, M. Théry, and K. John. 2019. Lattice defects induce microtubule self-renewal. Nat. Phys. 15:830–838. doi: 10.1038/s41567-019-0542-4.

Schatten, G. 1994. The Centrosome and Its Mode of Inheritance: The Reduction of the Centrosome during Gametogenesis and Its Restoration during Fertilization. Dev. Biol. 165:299–335. doi: 10.1006/dbio.1994.1256.

Schuh, M., and J. Ellenberg. 2007. Self-Organization of MTOCs Replaces Centrosome Function during Acentrosomal Spindle Assembly in Live Mouse Oocytes. Cell. 130:484–498. doi: 10.1016/j.cell.2007.06.025.

Sluder, G., F.J. Miller, and K. Lewis. 1993. Centrosome inheritance in starfish zygotes II: Selective suppression of the maternal centrosome during meiosis. Dev. Biol. 155:58–67. doi: 10.1006/dbio.1993.1006.

Srayko, M., E.T. O’Toole, A.A. Hyman, and T. Müller-Reichert. 2006. Katanin Disrupts the Microtubule Lattice and Increases Polymer Number in C. elegans Meiosis. Curr. Biol. 16:1944–1949. doi: 10.1016/j.cub.2006.08.029.

Staub, J., T.G. Setty, N.T. Nguyen, A. Paoletti, and P.T. Tran. 2005. Ase1p Organizes Antiparallel Microtubule Arrays during Interphase and Mitosis in Fission Yeast □. 16:1756–1768. doi: 10.1091/mbc.E04.

Syrovatkina, V., C. Fu, and P.T. Tran. 2013. Antagonistic spindle motors and MAPs regulate metaphase spindle length and chromosome segregation. Curr. Biol. 23:2423–2429. doi: 10.1016/j.cub.2013.10.023.

Szollosi, D., P. Calarco, and R.P. Donahue. 1972. Absence of centrioles in the first and second meiotic spindles of mouse oocytes. J. Cell Sci. 11:521–541.

Tanaka, K., and A. Hirata. 1982. Ascospore development in the fission yeasts Schizosaccharomyces pombe and S. japonicus. J. Cell Sci. Vol. 56:263–279.

Tanaka, T.U., M.J.R. Stark, and K. Tanaka. 2005. Kinetochore capture and bi-orientation on the mitotic spindle. Nat. Rev. Mol. Cell Biol. 6:929–942. doi: 10.1038/nrm1764.

Theurkauf, W.E., and R.S. Hawley. 1992. Meiotic spindle assembly in Drosophila females: Behavior of nonexchange chromosomes and the effects of mutations in the nod kinesin-like protein. J. Cell Biol. 116:1167–1180. doi: 10.1083/jcb.116.5.1167.

Tomita, K., and J.P. Cooper. 2007. The Telomere Bouquet Controls the Meiotic Spindle. Cell. 130:113–126. doi: 10.1016/j.cell.2007.05.024.

Unsworth, A., H. Masuda, S. Dhut, and T. Toda. 2008. Fission Yeast Kinesin-8 Klp5 and Klp6 Are Interdependent for Mitotic Nuclear Retention and Required for Proper Microtubule Dynamics. 19:5104–5115. doi: 10.1091/mbc.E08.

Vardy, L. 2000. The fission yeast gamma-tubulin complex is required in G1 phase and is a component of the spindle assembly checkpoint. EMBO J. 19:6098–6111. doi: 10.1093/emboj/19.22.6098.

Vukušić, K., R. Buđa, A. Bosilj, A. Milas, N. Pavin, and I.M. Tolić. 2017. Microtubule Sliding within the Bridging Fiber Pushes Kinetochore Fibers Apart to Segregate Chromosomes. Dev. Cell. 43:11–23.e6. doi: 10.1016/j.devcel.2017.09.010.

Walczak, C.E., and R. Heald. 2008. Mechanisms of Mitotic Spindle Assembly and Function. Int. Rev. Cytol. 265:111–158. doi: 10.1016/S0074-7696(07)65003-7.

Walczak, C.E., I. Vernos, T.J. Mitchison, E. Karsenti, and R. Heald. 1998. A model for the proposed roles of different microtubule-based motor proteins in establishing spindle bipolarity. Curr. Biol. 8:903–913. doi: 10.1016/S0960-9822(07)00370-3.

West, R.R., T. Malmstrom, and J.R. McIntosh. 2002. Kinesins klp5(+) and klp6(+) are required for normal chromosome movement in mitosis. J. Cell Sci. 115:931–940. doi: 10.1083/jcb.152.3.425.

West, R.R., T. Malmstrom, C.L. Troxell, and J.R. McIntosh. 2001. Two related kinesins, klp5+ and klp6+, foster microtubule disassembly and are required for meiosis in fission yeast. Mol. Biol. Cell. 12:3919–3932. doi: 10.1091/mbc.12.12.3919.

West, R.R., and J.R. McIntosh. 2008. Novel interactions of fission yeast kinesin 8 revealed through in vivo expression of truncation alleles. Cell Motil. Cytoskeleton. 65:626–640. doi: 10.1002/cm.20289.

Wolff, I.D., M. V. Tran, T.J. Mullen, A.M. Villeneuve, and S.M. Wignalla. 2016. Assembly of Caenorhabditis elegans acentrosomal spindles occurs without evident microtubuleorganizing centers and requires microtubule sorting by KLP-18/kinesin-12 and MESP-1. Mol. Biol. Cell. 27:3122–3131. doi: 10.1091/mbc.E16-05-0291.

Yamashita, A., M. Sato, A. Fujita, M. Yamamoto, and T. Toda. 2005. The Roles of Fission Yeast Ase1 in Mitotic Cell Division, Meiotic Nuclear Oscillation, and Cytokinesis Checkpoint. 16:1378–1395. doi: 10.1091/mbc.E04-10-0859.

Yan, H., and M.K. Balasubramanian. 2012. Meiotic actin rings are essential for proper sporulation in fission yeast. doi: 10.1242/jcs.091561.

Yoshida, M., and S. Sazer. 2004. Nucleocytoplasmic transport and nuclear envelope integrity in the fission yeast Schizosaccharomyces pombe. Methods. 33:226–238. doi: 10.1016/j.ymeth.2003.11.018.

Yukawa, M., C. Ikebe, and T. Toda. 2015. The Msd1-Wdr8-Pkl1 complex anchors microtubule minus ends to fission yeast spindle pole bodies. J. Cell Biol. 209:549–562. doi: 10.1083/jcb.201412111.

Yukawa, M., T. Kawakami, M. Okazaki, K. Kume, N.H. Tang, and T. Toda. 2017. A microtubule polymerase cooperates with the kinesin-6 motor and a microtubule cross-linker to promote bipolar spindle assembly in the absence of kinesin-5 and kinesin-14 in fission yeast. Mol. Biol. Cell. 28:3647–3659. doi: 10.1091/mbc.E17-08-0497.

